# Spatiotemporal visualization of DNA replication by click chemistry reveals bubbling of viral DNA in virion formation

**DOI:** 10.1101/2024.01.16.575809

**Authors:** Alfonso Gomez-Gonzalez, Patricia Burkhardt, Michael Bauer, Morten O. Loehr, Nathan W. Luedtke, Urs F. Greber

**Affiliations:** Department of Molecular Life Sciences, University of Zurich, Zurich, Switzerland; Department of Chemistry, McGill University, Montréal, Québec, Canada

**Keywords:** Virus assembly and infection, DNA replication, Liquid-liquid phase separation, Double-color click chemistry, Alkyne-tetrazine inverse electron-demand Diels-Alder ligation, Copper(I)-catalyzed azide-alkyne cycloaddition, Coacervates, Fluorescence microscopy

## Abstract

The organisation of human chromosomes reversibly changes in cell division, and irreversibly in apoptosis or erythropoiesis by DNA condensation and fragmentation processes. Yet, how viral replication in the nucleus affects host and viral chromatin organisation remains poorly understood. Here we used dual-color click chemistry to image human adenovirus DNA replication, demonstrating host chromatin compaction during active expansion of the viral replication compartment (VRC). Early-replicated viral DNA (vDNA) segregated from VRC and lost phospho-serine5-RNA Pol-II and viral DNA-binding protein (DBP), while late-replicated vDNA retained active RNA Pol-II, besides viral RNA-splicing and DNA-packaging proteins. Depending on replication and the assembly protein 52K, the late-stage VRCs gave rise to progeny by droplet formation of vDNA with GFP-tagged virion protein V into 52K biomolecular condensates. The study reveals distinct functions of early and late-replicated vDNA and provides insight into active and passive liquid phase separated zones conducive to selective genome packaging of nascent virions.

## Introduction

The nucleus compartmentalizes DNA, and constitutes a hub for transcription, replication, DNA editing, repair, as well as chromatin assembly, condensation and decondensation (*1*). The processes of cell division, apoptosis, spermatogenesis and erythropoiesis are testimony to the dynamic nature of eukaryotic chromosomes (*2-5*). Chromatin dynamics can be addressed through static and live cell imaging including high-speed super-resolution fluorescence microscopy (*6, 7*), cryogenic electron tomography (*8, 9*), and increasingly deep learning, artificial neural networks and computer vision procedures (*10-12*). For example, *in situ* imaging has provided crucial information on transcription initiation, elongation and processing (*13*), activation of origins in DNA replication (*14*), chemical modifications of DNA and associated histones in epigenetic activation and silencing (*15*), DNA repair (*16*), chromatin dynamics, or nuclear envelope breakdown and reformation in mitosis (*17*).

Alone, the nucleus not only controls gene activity and function in healthy cells, but it also is a hot spot for pathogen subversion, in particular DNA tumor viruses, retroviruses or negative-strand RNA viruses replicating in the nucleus (*18, 19*). DNA viruses, such as adenovirus (AdV) and herpes virus compete with the host for nucleotides, and block the synthesis of cell DNA while promoting viral DNA synthesis, as visualized by bioorthogonal chemistry (*20-22*). Ongoing viral DNA replication gives rise to DNA factories in the nucleus, so-called viral replication compartments (VRC). Yet, despite progress in imaging virus infections and rapid advance in AI-enhanced image analyses (*23-29*), our knowledge about dynamics in the functional organisation of infected nuclei has remained incomplete.

Here, we developed a dual-color visualization approach for replicating cellular and viral DNA (vDNA) of AdV-C5 and herpes simplex virus (HSV)-type 1 replicating in the cell nucleus. In humans, AdV-C5 causes self-limiting disease of the respiratory organs and disseminates to immune cells and mucosal tissue of the gastrointestinal tract (*30, 31*). HSV-1 infections can be asymptomatic, mild or life-threatening, and they last for the life-time of the individual due to viral latency in neurons and sporadic dissemination upon reactivation (*32*). Both viruses are used as vectors in clinical gene therapy (*33, 34*). They deliver a proteinaceous capsid with double-stranded DNA into the cytosol, engage microtubule-dependent motors for nuclear targeting, dock to the nuclear pore complex (NPC), uncoat and deliver their genome into the nucleus (reviewed in *35, 36, 37*). Both single AdV and HSV genomes can be tracked with copper(I) azide−alkyne cycloadditions (CuAAC), or in case of AdV, also immunostaining of the vDNA condensing protein VII (*20, 38-40*).

Viral early transcription activation goes along with remodeling of the viral chromatin, along with S-phase progression of the cell and inhibition of host DNA synthesis (*41-43*). Consistently, the AdV DNA is remodelled by deposition of histone H3.3, a histone variant added to DNA independently of replication (*43, 44*). AdV and HSV genomes persist in nuclei of immune cells or neurons where they express low or undetectable immediate early E1A and ICP8 proteins, respectively (*45-47*). Upon derepression of early gene expression by immuno-suppressive treatment or ectopic viral infection, however, AdV replicate and give rise to life-threatening viremia (*30, 48*).

Three viral proteins directly promote the formation of the AdV replication centers: DNA polymerase (Pol), single-strand DNA binding protein (DBP also known as 72K), and terminal protein (TP), which is synthesized as a precursor protein (p) and serves as a polymerase primer on the 5’ ends of the linear viral genome. VRCs grow radially throughout infection, and give rise to a large, yet unexplained excess of genomes over progeny particles (*23, 49*). Progeny particles are assembled after intermediate and late viral gene expression, the latter yielding a primary transcript of the major late transcription unit that is differentially spliced and polyadenylated resulting in L1 to L5 mRNAs (*50*). Intermediate-expressed proteins mediate vDNA packaging (IVa2, 22K, 33K, pIIIa), and scaffolding (52K), while late proteins make up the capsid (hexon, penton base, IIIa, fiber, VI, VIII, IX, L3-p23/protease) and the DNA-core (IVa2, V, VII, X, TP). Packaged capsids are proteolytically processed into infectious mature virions (*51*). Here we use novel combination of click chemistry and quantitative fluorescence microscopy to address the dynamics of replicating vDNA and vDNA packaging into nascent virions. Our results provide evidence supporting concurrent DNA and protein assembly for AdV-C5 particle formation.

## Results

### Vinyl-modified nucleosides are incorporated into AdV-C5 and HSV-1 genomes and are clickable in replication and entry

Bioorthogonal click-chemistry (clicking) allows biosynthetic tagging of proteins, lipids or nucleic acids with a wide range of functionalized molecules, including fluorophores. Unlike protein labelling, DNA labelling has, however, been limited to just a few click reactions in cells, including strain promoted azide-alkyne cycloaddition and inverse electron-demand Diels–Alder (IEDDA) (*52*). We tested if vinyl-2’-deoxy-nucleosides were incorporated into viral DNA during replication using conventional IEDDA bioorthogonal labelling by substituting the bulky trans-cyclo-octane used previously with a small vinyl-tag present in 5-vinyl-2’-deoxyuridine (VdU) (*53, 54*). VdU reacts with acridine orange-coupled 6-methyl-tetrazine (AO-6MT) (Fig. 1A). AO-6MT is a cell permeable, double-strand DNA intercalator, which increases the IEDDA reaction rate on DNA 60’000-fold compared to non-templated reactions (*54*). Human lung epithelial A549 cells were infected with AdV at high multiplicity of infection (MOI) for 24 h, incubated with VdU for 4 h, fixed, labeled with AO-6MT and stained for the viral replication compartment with DBP antibodies. VdU was incorporated into replicating vDNA (DBP-positive) in a concentration-dependent manner yielding flower-shaped patterns characteristic of VRC (Fig. 1B, C), akin to 5-ethynyl-2’-deoxycytidine (EdC) tagged and CuAAC-stained vDNA (*20*). Comparison of different vinyl-nucleosides showed that VdU was superior to 5-vinyl-2’-deoxycytidine (VdC) and 5-vinyl-2’-deoxyadenosine (VdA) in AdV as well as in HSV-1 infected cells, the latter co-stained with the HSV-1 replication marker ICP0 (Fig. S1A-C). Likewise, AO-6MT was superior to XFD488-6MT and 5-TAMRA-6MT, which did not react with VdU and thus gave no specific signal (Fig. S1D). Strikingly, VdU unlike EdC was incorporated into vDNA of E1-expressing HEK-293 and HER-911 cells infected with AdV-C5 (Fig. 1D, E, and Fig. S1E, F), and could be identified in CsCl-purified AdV-C5-VdU and E1-deleted AdV-C5-VdU (AdV-C5_ΔE1-VdU) particles by AO-6MT virion labeling, most notably independent of capsid disruption suggesting that AO-6MT penetrates into intact virions (Fig. 1F, G, and Fig. S1G, H). VdU-tagged AdV-C5 particles could be tracked in cell entry as indicated by immuno-stained capsids colocalizing with AO-6MT click-labeled vDNA 30 min pi, and vDNA dissociated from capsid 180 min pi (Fig. 1H). Strikingly, while AdV-C5 progeny formation was only slightly affected by VdU at concentrations up to 50 µM, HSV-1 titers were strongly reduced by low micromolar concentrations of VdU or VdC and yielded low amounts of HSV-1-VdU particles trackable by AO-6MT in A549 cell entry (Fig. S1I-L). Together, the data show that VdU is readily incorporated into AdV-C5 and less so into HSV-1 genomes, likely reflecting differences in the respective viral DNA polymerases.

**Figure 1:**
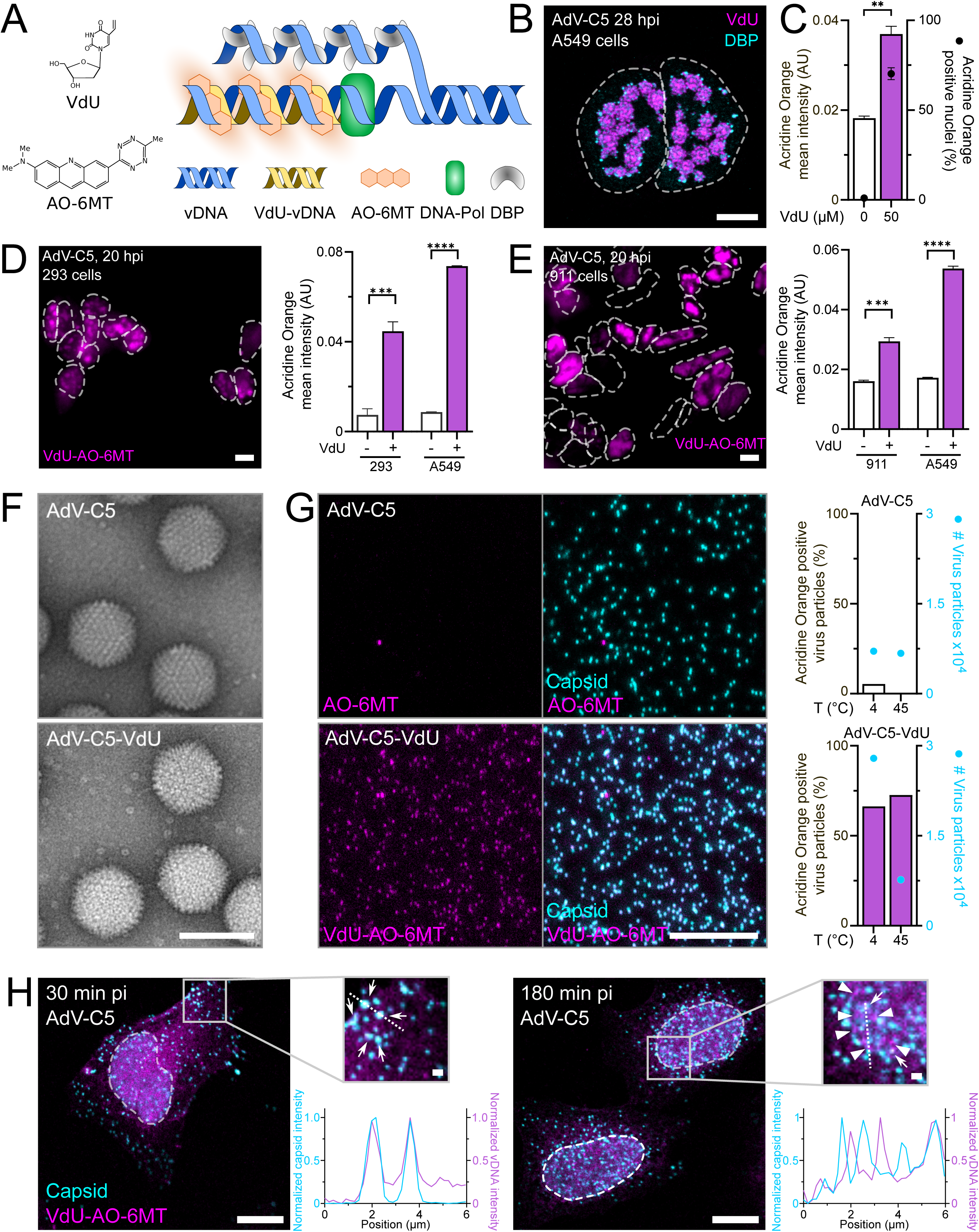
VdU is incorporated into replicated vDNA visualized throughout infection. (A) Tagging of newly synthesized 5’-vinyl-2’-deoxyuridine (VdU) modified viral DNA with Acridine Orange 6-methyl tetrazine (AO-6MT). (B) VdU incorporation during AdV-C5 replication. A549 cells where inoculated with AdV (MOI 3) at 37°C for 60 min, washed and fixed at 28 hpi. VdU was added 4h prior to fixation. Samples were immuno-stained for DBP and clicked with AO-6MT. Scale bar, 10 µm. (C) Quantitative analyses of VdU incorporation in A549 cells infected with AdV-C5 (MOI 1.5) and clicked with AO-6MT as described in B). Nuclei were segmented based on DAPI nuclear signal and the Acridine Orange signal was quantitated in the nuclear area. Data represent means ± SD. (D,E) VdU incorporation into AdV-C5 infected (MOI 1.5) HEK-293 and human embryonic retina (HER) 911 cells, fixed 20 hpi. For non-infected cells, VdU was added throughout 20 h, and for AdV-C5 infected cells 4 h prior to fixation. Samples were clicked with AO-6MT. Dashed lines indicate nuclear outlines. Scale bar, 10 µm. Bar graphs show quantitative analyses of VdU incorporation into HEK-293 and HER-911 cells. Data represent means ± SD of the AO-6MT signal over the DAPI-stained nuclei. Statistical significance was determined by non-parametric ANOVA with Holm-Sidak for multiple comparisons. ***, p<0.0002; ****, p<0.0001. (F) Electron micrographs of negatively stained AdV-C5 and AdV-C5-VdU particles. Scale bar, 100 nm. (G) Representative images and quantitative analysis of AdV-C5 and AdV-C5-VdU particles stained with AO-6MT. Virions were bound to poly-lysine-coated coverslips, fixed, stained with the anti-hexon 9C12 antibody and clicked with AO-6MT. Data represent means of the AO-6MT signal in virus particles. Scale bar, 10 µm. (H) Tracking of incoming VdU-labeled virions in cells. HeLa cells were inoculated with AdV-C5-VdU for 30 min, washed, incubated for 0 or 150 min, fixed, stained with anti-hexon 9C12 antibodies and clicked with AO-6MT. Arrows indicate VdU-AO-6MT labeled vDNA-containing viral particles; arrowheads indicate released VdU-AO-6MT vDNA. Data represent normalized intensities of capsid and vDNA across the dotted lines. Scale bar, 10 µm.

### Margination of viral and preexisting cellular chromatin during AdV replication

To explore the dynamics of cellular and AdV DNA in the course of infection, we developed a dual label approach using EdC and VdU pulses for CuAAC and IEDDA, respectively. We first asked if there was functional heterogeneity in the replicated vDNA over time. Pulse labeling of infected cells with EdC showed that vDNA synthesized after 16 hpi was incorporated into progeny virions harvested at 48 hpi, ranging from about 20% to nearly 80% of the purified virions containing detectable EdC from 16-24 h and 24-48 hpi pulsing, respectively (Fig. 2A). We then tested if EdC and VdU incorporation into VRC was comparable by using pulse labeling of A549 cells with EdC 16-20 hpi, followed by VdU 20-24 hpi or *vice versa*, and staining with anti-DBP antibodies (cyan), AO-6MT (magenta) and N_3_-AlexaFluor647 (green) at 24 hpi (Fig. S2A, B). Qualitative analysis showed extensive overlap of EdC and VdU with the VRC marker DBP, indicating that both EdC and VdU tags were authentic VRC reporters.

**Figure 2:**
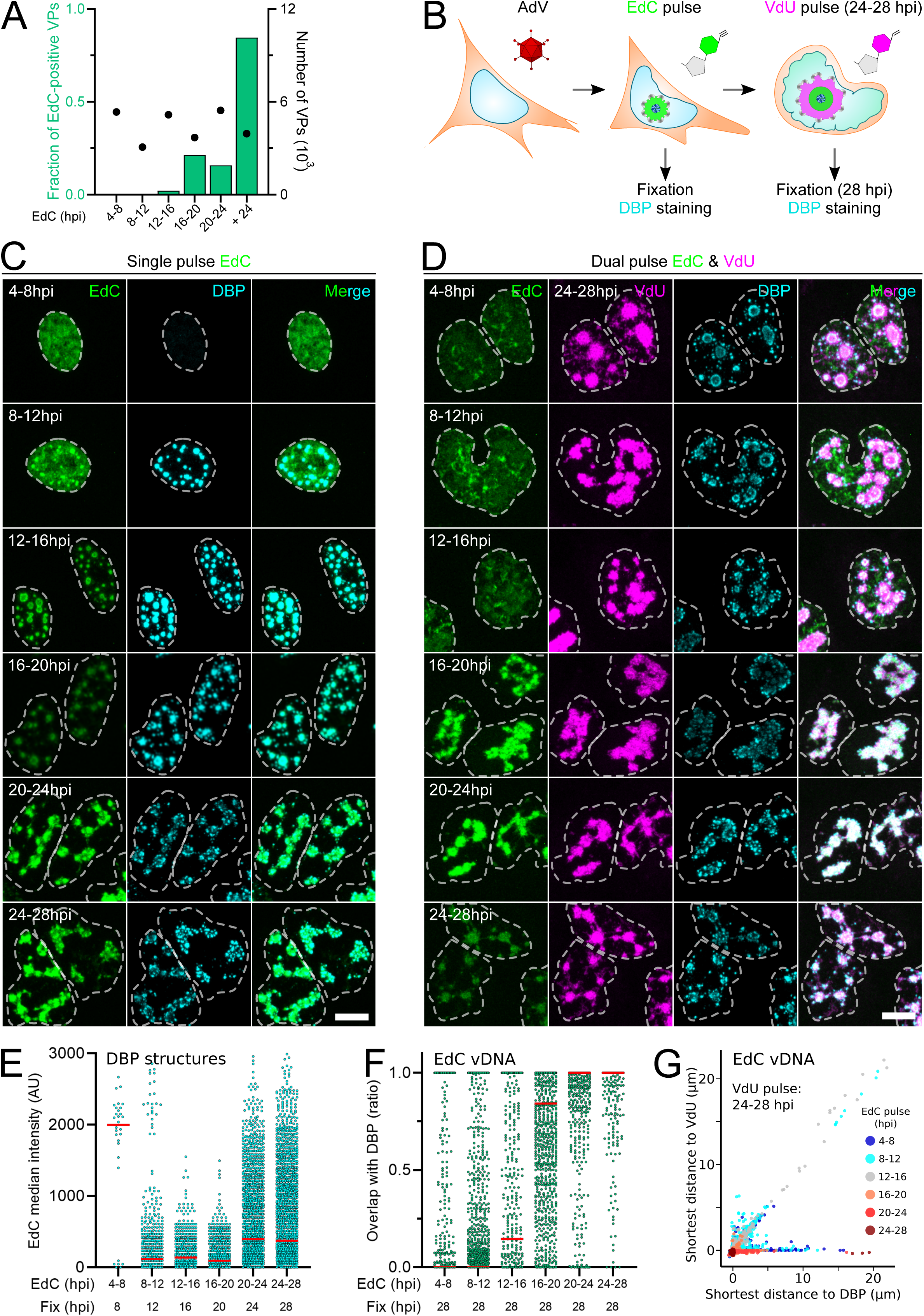
AdV-C5 replication centers evolve distinct structures and functions. (A) Encapsidation of differentially replicated viral DNA. Cesium chloride purified progeny virions from differentially EdC pulse-labeled A549 cells were heat disrupted, bound to poly-lysine-coated coverslips, fixed, anti-hexon 9C12 stained and N_3_-AlexaFluor488 clicked. Data represent the relative amount of or AlexaFluor488-positive capsid-stained virus particles. (B) Dual pulse-chase biorthogonal labeling of vDNA during AdV-C5 infection. A549 cells were infected with AdV-C5 (MOI 3) for 60 min, washed, pulse-labeled with 2.5 µM EdC for 4 h at different times pi, washed and either fixed, or tagged with 50 µM VdU at 24-28 hpi. Samples were stained with anti-DBP antibodies (cyan), clicked with AO-6MT (magenta) and clicked with N_3_-AlexaFluor647 (green). (C) AdV replication compartment visualized by EdC pulse labeling. Samples were prepared as indicated in B) (samples were fixed post EdC labeling). Images are maximum projections. Scale bar, 10 µm. (D) The VRC displays distinct patterns of early and late replicated viral DNA. Samples were prepared as indicated in panel B). Note that AO-6MT was added post fixation (after antibody staining) for 4 h at 37°C in the dark. Images are maximum projections. Scale bar, 10 µm. (E) Quantitative analysis of the EdC content in the VRC across infection. DBP structures from panel C) were 3D-segmented based on DBP signal and the EdC-AlexaFluor647 signal was quantitated in DBP volume. Each data point represent a DBP structure. Median is indicated in red. (F) Quantitative analysis of the vDNA dissociation from DBP across infection. EdC-labeled vDNA structures from panel D) were 3D-segmented based on EdC-AlexaFluor647 signal. Overlap was computed against the 3D-segmented DBP structures. Each data point represent an EdC-labeled vDNA structure. Median is indicated in red. (G) Quantitative analysis of the early-replicated vDNA distance to the actively replicating vDNA. EdC structures from panel D) were segmented as indicated in F). Data represents the shortest distance from single 3D-segmented EdC-labeled vDNA surfaces to both DBP and VdU-labeled vDNA structures within DAPI-stained nuclei, color-coded by the EdC pulse time.

The temporal dynamics of the VRC AdV-infected A549 cells were studied by EdC pulses for 4 h at different time points pi. As expected, the early EdC pulse (4-8 hpi) revealed a fairly homogeneous nuclear pattern lacking DBP signal, suggestive of cellular DNA in S-phase induced by the AdV early protein E1A (*55*) (Fig. 2B-D). When the EdC pulse was applied later in infection, the pattern gradually changed to discrete compartments that all became positive for VRC marker DBP at 20 hpi, and exhibited a noticeable increase in the mean EdC intensity at 24 hpi, indicating that VRC is highly dynamic (Fig. 2E, and Fig. S2C).

To further assess the spatiotemporal dynamics of the VRC, we used dual-label click experiments pulsing early replicated vDNA with EdC for 4 h, and late compartments by a VdU pulse right before chemical fixation at 28 hpi, followed by EdC-N_3_-AlexaFluor647 and VdU-AO-6MT staining (Fig. 2D). While volumetric analysis indicated an extensive overlap between VdU and the VRC marker and *vice versa* (Fig. S2D, E), the experiment revealed several striking features. One is that DNA synthesized before 16 hpi increasingly overlapped with DBP indicative of increasing vDNA, two the brightest spots of EdC (signal in green) segregated from the VRC stained with VdU-AO-6MT 24-28 hpi (pink signal), and three, the EdC signal progressively overlapped with VdU up to 24 hpi, as indicated by time-resolved shortest distance plots (Fig. 2F, G).

Similar to early-synthesized vDNA, cellular chromatin tagged with VdU prior to infection segregated from the VRC tagged with EdC (Fig. S2F, G). Together with live cell experiments where cellular DNA was labeled with VdU-AO-6MT prior to infection, these results demonstrated that viral DNA synthesis is not stalled and vDNA incorporates EdC in the presence of VdU-AO-6MT labeled cellular DNA (Fig. S2H, I, and Movie 1). The data also demonstrate the clustering of preexisting cellular DNA as a consequence of AdV-C5 infection, a notion which is underscored by the observation that DAPI-stained DNA condensed in the nuclear periphery as the nuclei extend in size (Fig. S2J).

Finally, we tested the possibility to image replicating AdV DNA by VdU-AO-6MT in live cells. A549 cells were infected with AdV-C5 (MOI 3), pulsed with VdU 18-22 hpi and subsequently labelled with the cell-permeable AO-6MT dye for the duration of the experiment. While the AO-6MT signal in the nuclei appeared to increase over time, its pattern remained static suggesting stalled replication, a notion confirmed by tagging the cells with EdC 24-28 hpi, fixation, and labeling with N_3_-AlexaFluor647 showing no double-positive AO-6MT / N_3_-AlexaFluor647 cells (Fig. S2K, L, and Movie 2). A similar result was obtained from HSV-1 infected cells pulsed with VdU 1-6 hpi, AO-6MT addition at 7 hpi and EdC tagging 12-16 hpi (Fig. S2M, N, and Movie 3). Collectively, the data show that VdU *per se* is not toxic to cellular or viral replication if used with short pulses of AO-6MT, although prolonged treatment with AO-6MT stalls DNA replication likely due to the bulky nature of the covalent DNA adduct (see Fig. 1A).

### Early-replicated vDNA has marks and localizations distinct from late-replicated vDNA

We took advantage of the dual-label click chemistry protocol to ask if early- and late-replicated vDNA differed with respect phosphorylated serine5-RNA Pol-II (p-RNA Pol-II), which indicates active transcription (*13*). Quantitative fluorescence microscopy showed that uninfected cells had a rather homogeneous p-RNA Pol-II pattern, while AdV-C5 infected cells strongly enriched the marker in the VRCs early in replication (14 hpi), as revealed by a single EdC pulse 12-14 hpi (Fig. 3A, B). When infection progressed to late stages e.g. 28 hpi, the early replicated vDNA partially segregated away from the late replicated p-RNA Pol-II-positive VdU-tagged vDNA in the VRC, and was p-RNA Pol-II-negative, as shown by quantification of total confocal projections (Fig. 3C). Intriguingly, the loss of the p-RNA Pol-II activation marker from the early replicated vDNA was strongly reduced by the DNA replication inhibitor 3′-deoxy-3′-fluorothymidine (DFT), which potently blocks AdV infection (*56*), as indicated by the blunted incorporation of VdU in vDNA (Fig. 3D). DFT blocked the expansion of the VRC and led to clustering of DBP, which largely colocalized with the early-replicated vDNA (Fig. 3E). Together, these results show that the early-replicated vDNA, which is not incorporated into progeny virions (see Fig. 2A), is transcriptionally active early in infection, but segregated from the VRC, and transcriptionally attenuated later on infection depending on vDNA replication.

**Figure 3:**
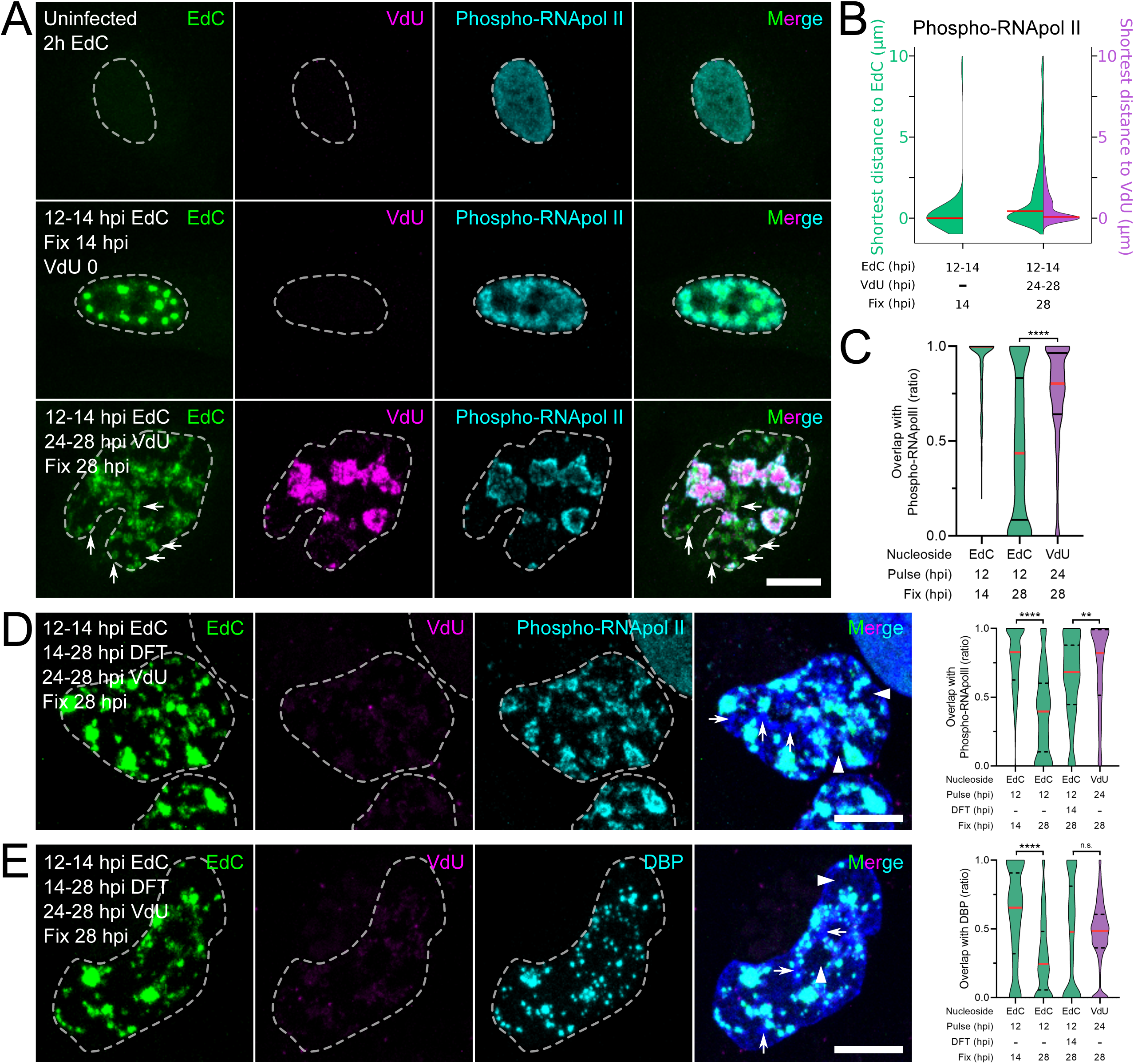
Phospho-RNA Pol-II is removed from early-replicated vDNA at late stage of infection but not late-replicated vDNA. (A) Distribution of p-RNA-Pol-II across AdV-C5 infected A549 cells. Specimens were labeled with 2.5 µM EdC at 12-14 hpi, EdC washed out and samples either fixed, or tagged with 50 µM VdU at 24-28 hpi, stained with anti-p-RNA Pol-II-CTD (cyan), and clicked with AO-6MT (magenta) and N_3_-AlexaFluor594 (green). Images are maximum projections. Scale bar, 10 µm. (B) Violin plots of quantitative analysis of the p-RNA Pol-II distance to the differentially pulse-labeled vDNA, as described in panel A). Data represent the shortest distance from single 3D-segmented p-RNA Pol-II structures to either EdC or VdU-labeled vDNA at different times pi. (C) Quantitative analysis of p-RNA Pol-II localization on early- and late-replicated vDNA pulsed with EdC for 2 h or VdU for 4 h, as described in A). Data represent co-localizing volumes of single EdC (green) and VdU (magenta) segmented 3D surfaces to p-RNA Pol-II within the DAPI-stained nuclei. Median is indicated in red. Statistical significance was determined by non-parametric ANOVA with Holm-Sidak for multiple comparisons. ****, p<0.0001. (D) Representative images and quantitative analysis of replication-dependent vDNA dissociation from active transcription sites. Samples were infected and labeled as described in A). DNA replication was inhibited with 2.5 µM 3′-deoxy-3′-fuorothymidine (DFT) at 14-28 hpi. Samples were stained with p-RNA Pol-II (cyan), and clicked with AO-6MT (magenta) and N_3_-AlexaFluor594 (green). Data distributions are shown as violin plots. Median is indicated in red. Statistical significance was determined by non-parametric ANOVA with Holm-Sidak for multiple comparisons. **, p<0.0021; ****, p<0.0001. Arrows indicate chromatin-dense nuclear regions; arrowheads indicate chromatin-free nuclear regions. Images are maximum projections. Scale bar, 10 µm. (E) Representative images and quantitative analysis of replication-dependent vDNA dissociation from the VRC. Samples were prepared as described in panel D). Samples were stained with anti-DBP (cyan), and clicked with AO-6MT (magenta) and N_3_-AlexaFluor594 (green). Statistical analyses was performed as indicated in panel D). p<0.0002; ****, p<0.0001; ns, non-significant. Arrows indicate chromatin-dense nuclear regions; arrowheads indicate chromatin-free nuclear regions. Images are maximum projections. Scale bar, 10 µm.

The formation of AdV progeny in the nucleus requires ongoing vDNA replication and synthesis of structural proteins from alternatively spliced late (L) mRNAs (L1-L4) driven by the major late promoter, which involves transactivation by the intermediate proteins L4-IVa2 and L4-22K (*57-59*). We used immunofluorescence microscopy to map L1-IIIa, L1-52K, L4-IVa2, L4-33K, L2-V and L2-VII with respect to early and late-replicated vDNA. As expected, the condensate-associated IIIa and 52K were homogenously distributed around the early and late vDNA in the nucleus without strong overlaps (although IIIa was also found also in the cytoplasm) (Fig. S3A, B), consistent with the notion that IIIa / 52K form biomolecular condensates around the VRC (*60*). In contrast, IVa2 exhibited distinct patterns in the nucleus. It is known to cooperatively bind to the vDNA packaging sequences for virion assembly, together with IIIa, 52K, 33K and 22K (*61-64*). Interestingly, about 25% of early vDNA overlapped with IVa2 at 14 hpi but only a few percent overlapped at 34 hpi, and about half of late-replicated vDNA was IVa2-positive at 34 hpi, suggesting that IVa2 preferentially marks packaging active vDNA (Fig. S3C). Similar to IVa2, 33K is involved in vDNA packaging, but also alternative splicing of late AdV transcripts (*65, 66*). It strongly overlapped with late- and to a lesser extent early-replicated vDNA suggesting that it is a marker of both early and late vDNA (Fig. S3D). Finally, we explored the location of the virion core proteins V and VII, which have capsid-vDNA bridging and condensing functions, respectively (*40, 67*). While V overlapped strongly with early- and late-replicated vDNA (Fig. S3E), a large fraction of late replicated vDNA and a smaller fraction of early-replicated vDNA were VII-positive at 34 hpi (Fig. S3F).

In sum, the data provide evidence for dynamic functions of replicating vDNA. Early-replicated vDNA is phospho-RNA Pol-II-positive early, and then segregates from VRC to become phospho-RNA Pol-II-negative depending on viral replication. At late stage of infection when virion assembly takes place, the large majority of early-replicated vDNA loses association with the vDNA packaging sequence-binding protein IVa2 while gaining association with 33K and VII, suggesting that this vDNA is no longer engaged in replication either. This notion is consistent with the lack of DBP on early-replicated vDNA (see Fig. 2D). In contrast, the late-replicated vDNA is largely positive for both IVa2 and VII, besides V, 33K and the condensate protein 52K, suggesting that it has an active role in virion morphogenesis.

### Formation of intermediate- and late-replicated IVa2-positive vDNA structures requires the assembly protein 52K

We analyzed the subnuclear localization of early- and intermediate-replicated vDNA pulse-labeled with EdC 12-14 and 22-24 hpi, respectively, compared to the DBP-positive VRC labeled with VdU at 32-34 hpi. While the early-replicated vDNA was segregated from the VRC, the intermediate vDNA was fully enclosed by peripheral VdU-AO-6MT and DBP signals, as indicated by volumetric reconstructions and quantitative analyses 34 hpi (Fig. 4A-D). In addition to the prominent VRCs, the infected nuclei invariably also contained distinct late-replicated VdU-AO-6MT vDNA puncta (bubbles) located between the VRC and the nuclear periphery 34 hpi (Fig. S3, marked with white squares). The puncta resembled coacervates, a synthetic polymer-, protein- or nucleic acid-dense aqueous phase formed by liquid-liquid phase separation (*68*). Strikingly similar puncta of intermediate-replicated vDNA pulse-tagged with EdC 22-24 hpi were observed between the VRC and the nuclear periphery next to VdU puncta pulsed 32-34 hpi, as quantified by shortest distance plots (Fig. 4E-G). All the intermediate and late vDNA puncta were DBP-negative (Fig. S4A, B). Together with our earlier observations that intermediate- and late-replicated vDNA is packaged into virions (Fig. 2), the results suggested that the punctate vDNA structures represent *bona fide* particle assembly intermediates. Interestingly, the puncta likely emerged from the periphery of the VRC, which stained positive for the vDNA packaging protein IVa2, although the puncta did not contain detectable amounts of IVa2 protein, possibly due to epitope masking or low levels of immunofluorescence signal (Fig. 4H, Fig. S4C). Likewise, all puncta were negative for IIIa and 52K, akin to the intermediate one lacking 33K, although the late-replicated ones were 33K positive in more than half of the cases (Fig. S4D-F).

**Figure 4:**
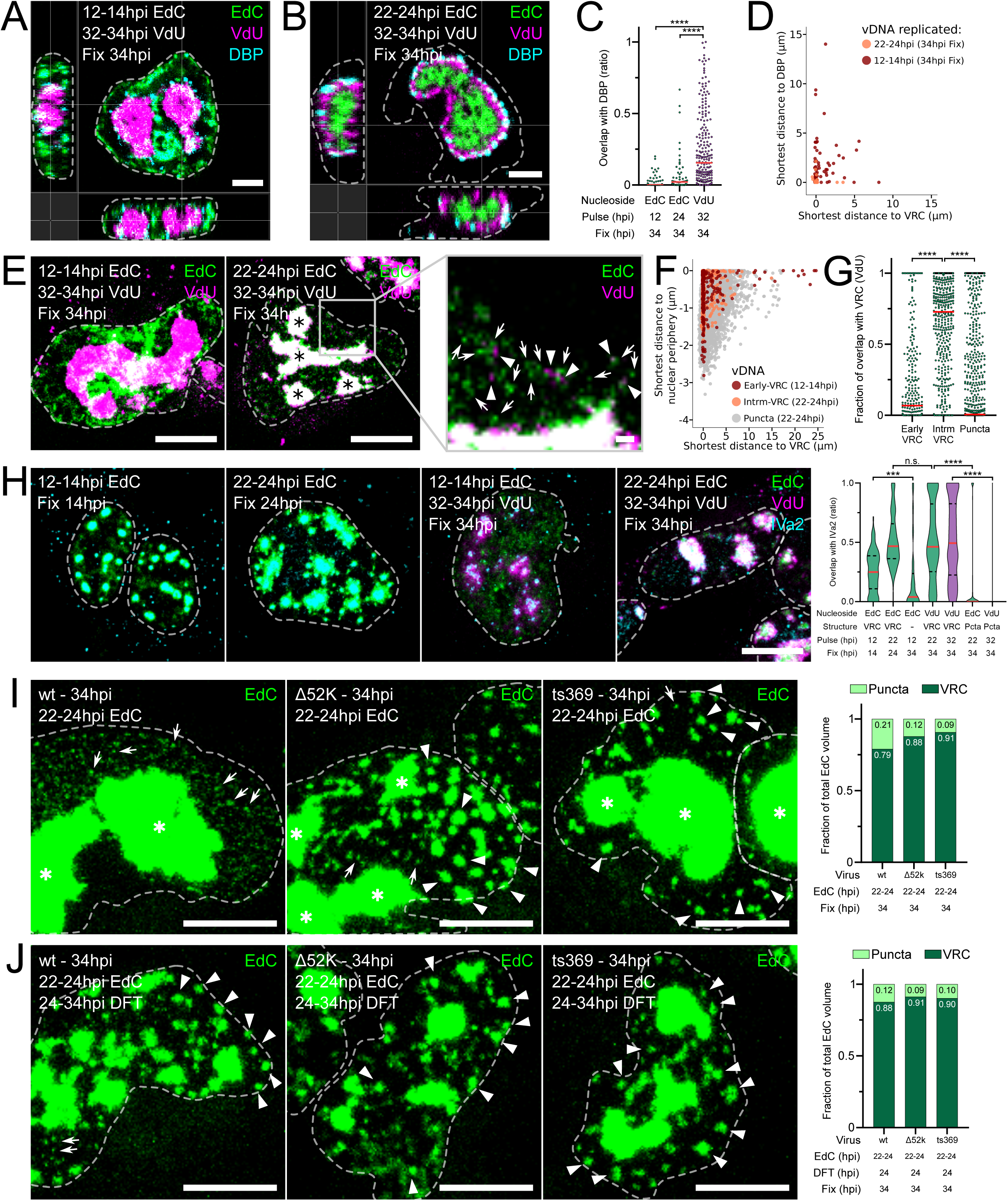
Late- but not early-replicated vDNA puncta structures dissociate from the VRC. (A,B) Early-replicated vDNA dissociates from the VRC, while the intermediate-replicated vDNA nucleates the mature VRC. Specimens were labeled with 2.5 µM EdC and 50 µM VdU as described in the image. Samples were stained with anti-DBP (cyan), and clicked with AO-6MT (magenta) and N_3_-AlexaFluor594 (green). Images are orthogonal slices. Scale bar, 5 µm. (C) Quantitative analysis of differentially-labeled vDNA co-localization with DBP at late stages of infection. Samples were prepared as in A) and B). EdC- and VdU-labeled vDNA structures were 3D-segmented based on the AlexaFluor594 and AO-6MT signals respectively. Overlap was computed against the 3D-segmented DBP structures. Each data point represents a vDNA structure. Median is indicated in red. Statistical significance was determined by non-parametric ANOVA with Holm-Sidak for multiple comparisons. ***, p<0.0002; ****, p<0.0001. (D) Quantitative analysis of differentially-labeled vDNA distance to the actively replicating vDNA at late stages of infection. Data represents the shortest distance from single 3D-segmented EdC-labeled vDNA surfaces to both DBP and VdU-labeled vDNA (VRC) structures within DAPI-stained nuclei, color-coded by the EdC pulse time. (E) Samples were prepared as described in A) and B). Black asterisks indicate VRCs; arrows indicate single EdC-labeled puncta (labeled at 22hpi); arrowheads indicate single VdU-labeled puncta (labeled at 32hpi). Images are maximum projections. Scale bar, 10 µm. Zoomed-in scale bar, 1 µm. (F) Quantitative analysis of the distribution of the different vDNA species across the nucleus. Samples were prepared as described in A) and B). Data represents the shortest distance from 3D-segmented EdC-labeled vDNA surfaces to both the nuclear periphery (based on the DAPI signal) and VdU-labeled vDNA (VRC) structures, color-coded by vDNA species. (G) Quantitative analysis of the different vDNA species co-localization with the most recently replicated vDNA (VRC) at late stages of infection. Samples were prepared as described in A) and B). Overlap was computed against the 3D-segmented VdU-labeled vDNA structures (VRC). Each data point represents an EdC-labeled vDNA structure. Median is indicated in red. Statistical significance was determined as indicated in C). ****, p<0.0001. (H) Representative images and quantitative analysis of time of synthesis-dependent vDNA co-localization with IVa2 during late stages of infection. Samples were prepared as described in A) and B). Data distributions are shown as violin plots. Median is indicated in red. Statistical significance was determined as indicated in C). ***, p<0.0002; ****, p<0.0001; ns, non-significant. Images are maximum projections. Scale bar, 10 µm. (I) Representative images and quantitative analysis of vDNA puncta in absence of a functional 52k protein. A549 cells were infected with either AdV-C5 wt, Δ52k or ts369 (at equivalent MOI), pulse-labeled with 2.5 µM EdC and clicked with N_3_-AlexaFluor594 as previously described. Data represents the proportion of puncta and VRC from the total EdC volume. White asterisks indicate VRCs; arrows indicate vDNA puncta; arrowheads indicate amorphous vDNA structures distinct from the VRC and puncta. Scale bar, 10 µm. (J) Representative images and quantitative analysis of vDNA puncta in absence of vDNA replication. Samples were prepared as described in I). DFT was added at 24 hpi and kept for the rest of the infection. Data represents the proportion of puncta and VRC from the total EdC volume. White asterisks indicate VRCs; arrows indicate vDNA puncta; arrowheads indicate amorphous vDNA structures distinct from the VRC and puncta. Scale bar, 10 µm.

Importantly, the EdC vDNA puncta around the VRC strongly depended on the presence of functional virion assembly protein 52K, as indicated by infections with AdV-C5 pm8001 lacking 52K (*69*), and AdV-C5_ts369 defective in infectious progeny formation at the restrictive temperature of 39.5°C (*70*) (Fig. 4I). Notably, both mutants did not affect VRC formation but appeared to increase VRC fragmentation. Likewise, the vDNA replication inhibitor DFT added right after the EdC pulse at 24 hpi impaired the formation of vDNA puncta in AdV-C5, pm8001 or ts369 infected cells at 34 hpi, besides fragmenting the VRC (Fig. 4J). Together the data suggest that the intermediate and the late-replicated vDNA structures are in the pathway of virion formation.

### GFP-V containing vDNA structures bubble off from VRC

Intermediate- and late-replicated vDNA give rise to puncta between the VRC and the nuclear periphery. To further characterize these structures, we used live cell imaging with a replication-competent green fluorescent protein (GFP)-V encoding virus (AdV-C2_GFP-V), which packages the GFP-V fusion protein into virions, albeit at reduced levels compared to AdV-C2 (*38*). As expected, GFP-V localized to the VRC labeled with EdC 22-24 hpi, and was largely accessible to anti-V antibody staining by immunofluorescence microscopy (Fig. 5A), akin to untagged protein V (Fig. S3). EdC puncta peripheral to the VRC, however, were strongly positive for GFP-V, but poorly accessible to anti-V antibodies, as indicated by gSTED superresolution microscopy and line scan analyses (Fig. 5A, B, and Fig. S5A, B). Live cell imaging at frame rates of 15 sec demonstrated that GFP-V puncta bubbled off from the VRC at high frequency and speed (Fig. 5C; and Movies 4, 5). In contrast, DFT-treatment not only drastically reduced the amount and motility of the bubbling GFP-V puncta, but also gave rise to apparent resorption of the rare puncta back into the VRC (Fig. S6C; and Movies 6, 7). These data suggest that ongoing DNA replication in the VRC is required for bubbling of GFP-V intermediate assembly structures from the VRC to the surrounding zone, and for allowing the intermediate structures to travel towards the nuclear periphery where fully assembled virus particles accumulate in the course of infection (Fig. 5D).

**Figure 5:**
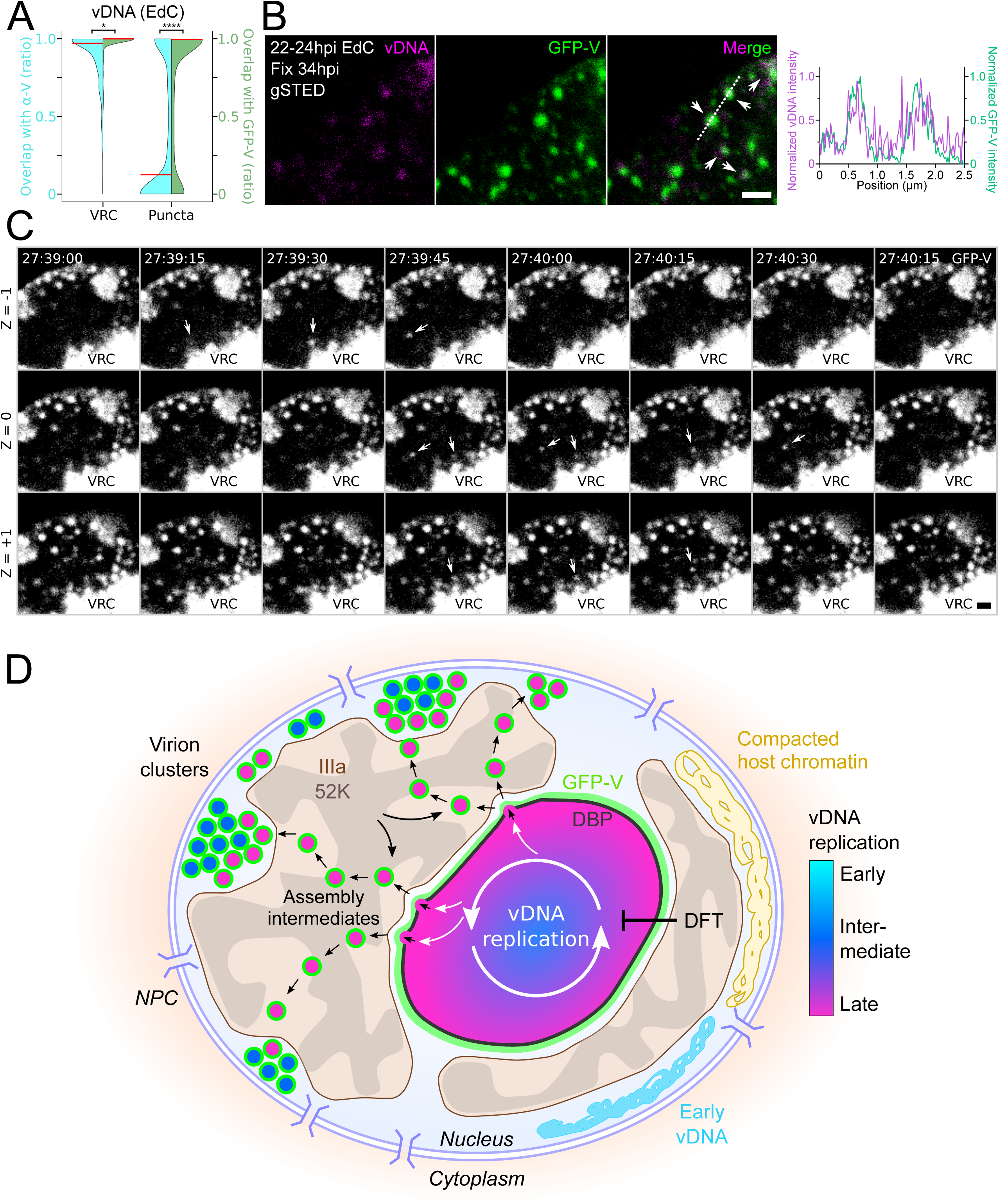
Punctate viral assembly intermediates containing vDNA and GFP-V bubble from VRC towards the nuclear periphery. (A) Viral DNA puncta are GFP-V positive but anti-V negative. A549 cells were infected with AdV-C2-GFP-V for 60min, washed, pulse-labeled with 2.5 µM EdC at 22-24 hpi, EdC was washed out and cells were fixed at 34 hpi. Samples were stained with anti-V and clicked with N_3_-AlexaFluor594. EdC-labeled vDNA structures were 3D-segmented based on the AlexaFluor594 signal and classified based on their size and intensity. Overlap was computed against the 3D-segmented GFP-V and anti-V antibody structures. Data distributions are shown as violin plots. Median is indicated in red. Statistical significance was determined by non-parametric ANOVA with Holm-Sidak for multiple comparisons. *, p<0.03; ****, p<0.0001. (B) Viral DNA puncta can be found within the GFP-V puncta through gSTED. Samples were prepared as in A). samples were clicked with N_3_-AlexaFluor647. Samples were imaged in a SP8 gSTED microscope. Arrows indicate EdC-labeled vDNA positive GFP-V puncta. Data represent normalized intensities of vDNA (magenta) and GFP-V (green) across a dotted line. Scale bar, 1 µm. (C) Live imaging of AdV-C5 viral core morphogenesis. A549 cells were infected with AdV-C2-GFP-V as described in A). GFP-V signal was recorded on full cell Z-stacks from 27:30-28:00 hpi at a frequency of four stacks per minute. Arrowheads indicate viral core nucleation events. Images shown are three contiguous Z planes. Scale bar, 1 µm. (D) Model for vDNA in AdV morphogenesis involving dynamic assembly intermediates emerging from viral replication centers, and depending on active viral DNA replication zones and 52K / IIIa condensates.

## Discussion

Visualization of nucleic acids at single molecule resolution has given deep insight into host cell and viral dynamics. Underlying approaches include fluorescent in situ hybridization (FISH) (*71*), azide−alkyne cycloadditions (*72*), and a range of fluorescent reporter proteins, including split GFP (*73*), SunTags (*74, 75*), LacO/LacR, TetO/TetR or orthogonal dCas9 requiring genetic knock-in of repetitive target sequences (*76*) (*77, 78*), or the so-called ANCHOR system derived from bacterial ParB proteins binding to either of two centromeric regions of bacterial DNA (*79*). While FISH allows for multiplexing at high sensitivity, especially when combined with branched DNA technology (*80*), fluorescent reporters are applicable to both chemically fixed and live cells. Their use has remained challenging, however, due to variability in target identification, overexpression, and multimerization of up to hundreds of fluorescent tags on a target sequence (*81*). This can be problematic, as even small tags, such as lysine-cysteine-lysine (KCK) affect target DNA compaction, for example in case of ParB and ParB binding to target DNA (*82*).

In contrast, direct labeling of nucleic acids by chemical metabolites, such as alkyne-modified nucleosides, provides distinct advantages, including low invasiveness and near quantitative bioorthogonal coupling (*53, 83, 84*). Here we developed a double-enhanced strategy for nucleic acid-templated reactions in virus-infected cells utilizing a fluorogenic DNA-intercalating agent AO-6MT capable of undergoing IEDDA reaction with VdU-tagged DNA at unprecedented sensitivity and live cell resolution. The VdU-AO-6MT-labeled vDNA was bright enough to allow tracking of vDNA entities in the periphery of VRC and in incoming single virus particles and it overcomes the narrow substrate specificity of endogenous nucleotide kinases in E1-transformed cells thereby allowing for the first time to click-label AdV vector DNA. Unlike a recently reported pronucleotide-triphosphate analogue linked to 2-trans-cyclooctene (*53*), our protocol is fully compatible with serum-containing medium and not affected by carboxyesterase or lipase degradation. Viral and host DNA labeled with VdU-AO-6MT can be observed by live cell imaging for several hours although it eventually stalls DNA replication and reduces signal to noise upon prolonged times, leading to apoptosis as reported in uninfected cells (*85*).

The combination of IEDDA-VdU with CuAAC in pulse-chase experiments showed that the early-replicated AdV DNA along with preexisting host DNA was compacted, and, upon prolonged infection, became devoid of both transcription (p-RNA Pol-II) and replication-active (DBP) marks. A fraction of compacted early-replicated vDNA was positive for VII consistent with the notion that VII condenses the Xenopus chromatin and inhibits transcription and replication (*86, 87*). Consistently, early-replicated vDNA was not incorporated into viral progeny, known to contain unmethylated DNA (*88*). Possibly, compacted vDNA is CpG-methylated, as reported with abortive AdV infections (*88*). In analogy with HSV-1 capsids, which exhibit restricted diffusion by compacted chromatin near the nuclear envelope (*89*) and depend on expanded host heterochromatin for nuclear egress (*90*), we suggest that host chromatin compaction provides a volume free of cellular chromatin around the AdV VRC, and thereby facilitates the diffusion of vDNA bubbles promoting viral morphogenesis.

How progeny AdV particles are formed has remained controversial, though. A long standing sequential model postulated encapsidation of viral genomes into empty capsids akin to bacteriophages, largely based on observations that AdV-C5 particles contain a packaging ATPase (IVa2), and that infection yields empty and partially filled capsids besides fully packaged ones (*91, 92*). An alternative view is emerging based on *in situ* electron microscopy evidence for semi-closed capsid assembly intermediates suggesting that capsid assembly and DNA encapsidation are concurrent processes (*93*). By employing quantitative time-resolved *in situ* tracking of newly synthesized DNA, we now show that late- but not early-replicated viral genomes bubble from the VRC with associated packaging and core proteins, and late-replicated vDNA is assembled into progeny particles, depending on DNA synthesis, and the 52K assembly protein. 52K forms biomolecular condensates, and functions as a scaffold by bringing the capsid together with the viral genome in nascent virions (*60, 94*).

Virus particle formation may thus start at the interface of two distinct nuclear domains, the VRC and the 52K condensate. By analogy with chemically-fuelled phase separated droplets (*95*), VRC can be considered a nonequilibrium condensate, akin to nucleoli, PML bodies or transcriptional condensates (*13, 96*), but unlike the quasi-equilibrated 52K condensate. The interface between these compartments may give rise to structures with emerging properties, such as the bubbles containing nascent virus particles. Besides vDNA, the AdV bubbles contained GFP-V and 33K, but not IVa2, DBP, 52K and IIIa, suggesting that these epitopes were not accessible, reminiscent of an anti-V impermeable nuclear compartment late in infection (*97*).

While the nature of the bubble motions is still unknown, it was possible to freeze the motions by DNA polymerase inhibitors suggesting that they stem from outward pushing forces of the VRC, possibly DNA polymerization. This would be consistent with the observation that active droplets position their passive particles in their center (*98*). Thus, the active regions at the periphery of the VRC would drive the liquid-to-solid transition observed as bubbling events (*99*). Another source of force for bubble movement could be the collapse of viral or cellular DNA outside the VRC, as this DNA transitions towards compaction. Compaction might exert pulling forces on the bubbles containing nascent virus particles, and may eventually result in the accumulation of complete virions near the nuclear periphery, as observed at late stage of AdV-2_GFP-V infection by light and electron microscopy (*38*). The bubbles themselves are not replication-active, yet their protein composition is dynamic, suggesting that they are an indermediate in virion assembly. Assembled virions in well-ordered paracrystalline nuclear super-assemblies are reminiscent of liquid crystals, and eventually disperse upon rupture of the nuclear membrane and the lytic release from the infected cell (*100, 101*). In sum, our study here provides evidence for selective genome packaging into viral particles at the interface between an active compartment releasing vDNA reaction products, and a passive compartment made up of the biomolecular condensate protein 52K for particle assembly.

## Materials and Methods

### Cell culture and virus production

HeLa-ATCC, A549, HEK293T, and HER911 cells were maintained in Dulbecco’s modified Eagle’s medium (DMEM; Gibco) supplemented with non-essential amino acids (Thermo Fisher Scientific) and 10% fetal calf serum (FCS; Gibco). During infection experiments, the medium was additionally supplemented with penicillin (100 U/ml) and streptomycin (100 µg/ml). The cells were grown at 37°C in a 5% CO_2_ atmosphere for no longer than 20 passages.

All AdVs were grown in A549 cells and purified over two cesium chloride gradients as previously described (*102*). AdV-C5 (wt300) has been previously described (*103*). AdV-C5_Δ52K and ts369 were provided by Dr. Michael Imperiale (University of Michigan Medical School, United States of America) (*69*) and David Ornelles (Wake Forest University, Winston-Salem, North Carolina, USA) (*70*). AdV-C2–GFP-V was used as described (*38*). VdU-labeled AdV-C5 was produced as previously described (*20*), with the following modifications: VdU was added to the cells at 4 hpi to a final concentration of 50 µM and kept until harvest and purification. HSV-1 strain F (VR-733) was purchased from ATCC. HSV-1 viruses were pelleted from the supernatant and purified over one Nycodenz® gradient (SERVA Electrophoresis GmbH, Heidelberg) as previously described (*39*).

### Electron microscopy

For the analysis of purified AdVs, 5 µl of glycerol-free purified virus was mounted onto carbon-coated grids for 5 min. Grids were washed three times with distilled water and stained for 30 s with 10 µl of a 2% aqueous uranyl acetate solution. Samples were imaged in a CM100 transmission electron microscope at 80 keV (Thermo Fisher Scientific, Eindhoven, The Netherlands).

### Antibodies

The following primary antibodies were used: Rabbit anti-AdV protein VI, IF 1:2000 (*104*). Mouse anti-AdV-C5 hexon 9C12, IF 1:40 (*105*). Mouse anti-AdV DBP, IF 1:150 (provided by A. Levine, *106*). Rabbit anti-AdV IVa2, IF 1:100 (provided by P. Hearing, *107*). Rabbit anti-AdV 33K, IF 1:250 (provided by P. Hearing, *62*). Rabbit anti-AdV IIIa, IF 1:1000 (provided by P. Hearing, *108*). Rabbit anti-AdV 52K, IF 1:1000 (provided by P. Hearing, *107*). Rabbit anti-AdV V, IF 1:100 (*38*). Rabbit anti-AdV VII, IF 1:100 (provided by U. Petterson). Rabbit anti-HSV HC, IF 1:1000 (provided by R. Eisenberg and G. Cohen, University of Pennsylvania, Philadelphia, USA). Mouse anti-HSV ICP8, IF 1:10 (HB-8180; ATCC, Rockville, Md.). Mouse anti-phospho-Ser5-RNA-Pol-II, IF 1:1000 (ab5408, Abcam).

### Virus titrations and MOI determination

Ten thousand cells were seeded per well in a black 96-well imaging plate. On the following day, viruses were diluted in infection medium (DMEM supplemented with 2% FCS, non-essential amino acids, and penicillin/streptomycin). The culture supernatant was aspirated, and cells were inoculated with 100 µl of virus dilution for 60 min at 37°C, subsequently inoculate was removed, and cells were supplemented with 100 µl of fresh infection medium. The cells were fixed at 24 hpi with 3% paraformaldehyde (PFA) in phosphate-buffered saline (PBS) for 15min at room temperature. The remaining PFA was quenched with 25 mM NH4Cl diluted in PBS for 10 min, followed by permeabilization with 0.5% Triton X-100 in PBS for 5min. Cells were stained with rabbit anti-protein VI (*104*) diluted in blocking buffer (10% goat serum in PBS) for 1h at 4°C. After three washes of 3min each in PBS, cells were stained with secondary antibody (goat anti-rabbit AlexaFluor488, Thermo Fisher Scientific) diluted in blocking buffer containing 4′,6-diamidino-2-phenylindole (DAPI; 1 µg/ml) at room temperature for 1 h. After three more washes of 4 min in PBS, cells were imaged in a Molecular Devices high-throughput microscope (IXMc) in wide-field mode with a 20× objective. For quantification of infection with CellProfiler (*109*), nuclei were segmented according to the DAPI signal, and the intensity of protein VI over the nuclear mask was measured. MOI was defined by the probability of infection based on AdV protein VI expression according to the adapted Poisson distribution by Ellis and Delbrück (*110*).

### Virus progeny production and TCID50 titration

Fifteen thousand cells were seeded per well in a transparent 96-well plate. On the following day, viruses were diluted in infection medium to a final MOI of 1.5. The cell culture supernatant was aspirated, and cells were inoculated with 100 µl of virus dilution for 120 min at 37°C, subsequently inoculate was removed, cells were supplemented with DMSO or vinyl-modified nucleosides diluted in fresh infection medium and kept until cytopathic effect was observed (∼48 hpi for HSV-1 and ∼72 hpi for AdV-C5). For HSV-1 progeny, supernatant was collected, debris was removed by centrifugation at 4000g for 3 min, and remaining supernatant was snap frozen in liquid nitrogen and stored at −80 °C until titration. For AdV-C5 progeny, supernatant and cells were collected, freeze-thawed twice in liquid nitrogen, snap frozen a third time and kept at -80°C until use; upon use samples were thawed a third time and debris was removed by centrifugation at 4000g for 3min.

For progeny titration ten thousand cells were seeded per well in a transparent 96-well plate. On the following day, the collected virus was diluted in infection medium, added on top of the cells, and kept for seven days at 37°C. Seven days post infection cells were fixed by adding 16% PFA on top to reach a final concentration of 3% for 30 min at room temperature. Fixative was discarded and cells were stained with 40 µl of crystal violet staining solution (2.5mg/ml crystal violet, 10% MeOH in ddH2O) for 60 min at room temperature. Excess staining was washed away multiple times in ddH2O, wells were scored as infected when the cell monolayer wasn’t intact, and TCID_50_ values were obtain according to the Spearman-Kärber method.

### IEDDA click chemistry and vDNA analysis

Eighty thousand A549 cells were seeded on 12 mm glass coverslips in a 24 well plate. On the following day, viruses were diluted in infection medium to a final MOI of 3. The cell culture supernant was aspirated, and cells were inoculated with 300 µl of virus dilution for 60 min at 37°C, subsequently inoculate was removed and fresh infection media was added to the cells. Infected cells were tagged with 50 µM VdU 4 h in 2% fetal calf serum-containing medium prior to fixation. Non-infected samples were labeled with 50 µM VdU for the entire duration of the experiment. Cells were fixed with 3% PFA for 15min at RT, quenched with 25 mM NH_4_Cl for 10 min, and permeabilized with 0.5% Triton X-100 for 5min. Cells were stained with mouse anti-DBP (provided by A. Levine) (*106*) for AdV-C5 or with mouse anti-ICP8 (HB-8180; ATCC, Rockville, Md.) for HSV-1 in blocking buffer for 1h at 4°C. Cells were washed three times 3min each in PBS, and stained with secondary antibody goat anti-mouse AlexaFuor594 (Thermo Fisher Scientific), unless specified, in blocking buffer for 1h at RT. After primary and secondary antibody incubation, the coverslips were inverted onto a 35-µl droplet of freshly prepared IEDDA click reaction mix (50 µM AO-6MT in PBS) for 4h at 37°C. Samples were stained with DAPI and imaged in a Leica SP8 FALCON cLSM. Three dimensional image stacks were analyzed using Imaris10 (Oxford instruments, Oxon, UK). VdU-AO-6MT structures were segmented within the DAPI delimited volume and classified into VRC and puncta based on a voxel size threshold of 50 (image voxel size: X, 0.18 µm; Y, 0.18 µm; Z, 0.5 µm). Images were batch-processed and segmentation was manually validated. Overlap and distances were computed between the volumes within the nucleus.

### CuAAC click chemistry and vDNA analysis

Eighty thousand A549 cells were seeded on 12 mm glass coverslips in a 24 well plate. On the following day, cells were infected as previously described. Infected cells were labeled with 2.5 µM EdC 4 h prior to fixation unless specified. Non-infected samples were labeled with 2.5 µM EdC for the entire duration of the experiment. Cells were fixed, quenched, and permeabilized as previously described. Cells were stained with mouse anti-DBP (B6-8; provided by A. Levine) for AdV-C5 or with mouse anti-ICP8 (HB-8180; ATCC, Rockville, Md.) for HSV-1 in blocking buffer for 1h at 4°C, and stained with secondary antibody goat anti-mouse AlexaFuor594 (Thermo Fisher Scientific), unless specified, in blocking buffer for 1h at RT. After primary and secondary antibody incubation, the coverslips were inverted onto a 30- µl droplet of freshly prepared CuAAC click reaction mix [10 µM N_3_-AlexaFluor647 (Thermo Fisher Scientific), 1 mM CuSO_4_, 10mM aminoguanidine (Sigma-Aldrich), 1 mM Tris(3-hydroxypropyltriazolylmethyl)amine (THPTA) (Sigma-Aldrich) and 10 mM sodium ascorbate in PBS] for 2h at RT. Samples were stained with DAPI and imaged in a Leica SP8 FALCON cLSM. Three dimensional image stacks were analyzed using Imaris10 (Oxford instruments, Oxon, UK). EdC-N_3_-AlexaFluor structures were segmented within the DAPI delimited volume and classified into VRC and puncta based on a voxel size threshold of 50 (image voxel size: X, 0.18 µm; Y, 0.18 µm; Z, 0.5 µm). Images were batch-processed and segmentation was manually validated. Overlap and distances were computed between the volumes within the nucleus.

### Virion analysis on glass coverslips

Glass coverslips were coated with a 120-µl drop of 0.1 mg/ml poly-L-Lysine (Sigma-Aldrich) for 30 min at RT. On the meantime, 0.3 micrograms of purified virus were diluted in HEPES buffer (25mM HEPES, 150mM NaCl, 1mM MgCl_2_ in ddH_2_O) and incubated for 12min at on ice or at 45°C in a waterbath. Excess poly-L-Lysine was removed, mock or heat-treated viruses were added on the poly-L-Lysine coated coverslips for 30 min at RT and fixed with 3% PFA for 15 min at RT. Samples were quenched with NH_4_Cl for 10 min, stained with mouse anti-hexon 9C12 (*105*) for AdV-C5 or with rabbit anti-HC (kindly provided by R. Eisenberg and G. Cohen, University of Pennsylvania, Philadelphia, USA) for HSV-1 in blocking buffer for 1 h at 4°C and stained with secondary antibody goat anti-mouse/rabbit AlexaFuor680 (Thermo Fisher Scientific) in blocking buffer at RT for 1h. Samples were clicked with either IEDDA or CuAAC reactions as described above. Coverslips were mounted onto slides in ProLong^TM^ Gold Antifade Mountant (Thermo Fisher Scientific), let dry over two nights and imaged in a Leica SP8 FALCON cLSM as described.

### Virus particle entry and vDNA release

Eighty thousand HeLa cells were seeded on 12 mm glass coverslips in a 24 well plate. On the following day, cells were incubated with ca. 100 bound particles of virus in infection medium for 30 min at 37°C as described in (*40*). Inocula were removed and cells were fixed or incubated in fresh infection medium for 150 min at 37°C. Cells were fixed, quenched and permeabilized as previously described. Cells were stained with mouse anti-hexon 9C12 for AdV-C5 or with rabbit anti-HC for HSV-1 in blocking buffer for 1h at 4°C, stained with secondary antibody goat anti-mouse/rabbit AlexaFuor680 (Thermo Fisher Scientific) in blocking buffer for 1h at RT and IEDDA-clicked as described above. Coverslips were mounted onto slides in ProLong^TM^ Gold Antifade Mountant, let dry over two nights and imaged in a Leica SP8 FALCON cLSM as described.

### Dual click chemistry

Eighty thousand A549 cells were seeded on coverslips. On the following day, viruses were diluted in infection medium to a final MOI of 3. The cell culture supernatant was aspirated, and cells were inoculated with 300 µl of virus dilution for 60 min at 37°C, subsequently inoculate was removed and fresh infection media was added to the cells. Infected cells were pulse-labeled with 2.5 µM EdC for 4h at the specified times, EdC was washed out two times with PBS and fresh infection medium was added. Cells were labeled with 50 µM VdU 4h prior to fixation unless specified. Cells were fixed, quenched and permeabilized as previously described. Samples were stained with mouse anti-DBP and secondary goat anti-mouse AlexaFluor594 as previously described unless specified. Samples were firstly IEDDA-clicked flowed-up by the CuAAC click reaction as previously described. Samples were stained with DAPI, mounted onto slides in ProLong^TM^ Gold Antifade Mountant and imaged in a Leica SP8 FALCON cLSM. Three dimensional image stacks were analyzed using Imaris10 as previously described.

### Confocal microscopy

A Leica SP8 FALCON or SP8 gSTED cLSM was used in all experiments, in which single viral genomes and subnuclear structures were imaged. Imaging was performed with a 63× magnification oil objective with a numerical aperture of 1.40 and a zoom factor of 2, with a pixel size of 0.181 µm. Z stacks were captured with a step size of 0.3-0.5 µm to capture the entire cell, and the size of the pinhole was 1 Airy unit. For the acquisition of virus particle on coverslips, the pinhole was 6.28 Airy units. Leica hybrid detectors were used for each channel.

### Confocal spinning-disk live microscopy

Ten thousand A549 cells were seeded in a black glass bottom 96-well imaging plate. On the following day, cells were inoculated with virus at a final MOI of 3 for 60 min at 37°C, subsequently inoculate was removed, and cells were supplemented with 100 µl of fresh infection medium. Non-infected cells were labeled with VdU upon seeding and washed before infection. Infected cells were pulse-labeled with VdU at 16-20 hpi for AdV-C5 or at 1-6 hpi for HSV-1. Subsequently, medium was removed and samples were clicked with AO-6MT in FluoroBrite (Thermo Fisher Scientific) containing Hoechst 33342 as indicated. Imaging started 1 h post AO-6MT addition at a frequency of two to four frames per hour in a microscope IXMc spinning disk microscope in confocal mode with a 20× objective and a 60 µm pinhole. Samples were fixed, quenched, permeabilized, stained and re-imaged as described above. Samples were quantified with CellProfiler as described above.

For the imaging of viral core bubbling events, fifteen thousand A549 cells were seeded in 8-well µ-slides (Ibidi) over two nights. Cells were inoculated with AdV-C2-GFP-V at a final MOI of 3 in 180 µl of infection media for 60min at 37°C, subsequently inoculate was removed, and cells were supplemented with 300 µl of fresh infection medium. Media was replaced with mock or 2.5 µM DFT supplemented FluoroBrite (Thermo Fisher Scientific) at 24 hpi. Imaging started at the specified times at a frequency of four full-cell Z-stacks per minute in an Olympus IXplore SpinSR10 spinning-disk microscope with a 100x objective in silicon oil.

### Superresolution gSTED microscopy

Eighty thousand A549 cells were seeded on coverslips. On the following day, AdV-C2-GFP-V was diluted in infection medium to a final MOI of 3. Cells were infected and pulse-labeled with EdC as described previously. Cells were fixed with 2% PFA for 15min at RT, quenched with 25 mM NH_4_Cl for 10 min, and permeabilized with 0.1% Triton X-100 for 10 min. Cells were blocked with blocking buffer at 4°C for 1 h. Samples were CuAAC-clicked with a N_3_- AlexaFluor647 as described previously. Coverslips were mounted onto slides in ProLong^TM^ Gold Antifade mounting medium, let dry over two nights and imaged in a Leica SP8 inverse STED 3X cLSM. Imaging was performed with a 100x magnification oil objective with numerical aperture of 1.4 and a zoom factor of 6, with a pixel size of 19 nm and gating from 1.5 to 6 ns. A 592 nm depletion laser was used for the GFP, a 775 nm depletion laser was used for N_3_-AlexaFluor647.

### Statistical analyses

All graphs were generated using RStudio or GraphPad Prism and display means ± SD unless stated otherwise. Statistical tests used are indicated in the figure legends (ns, not significant; *P < 0.05; **P < 0.01; and ***P < 0.001).

## Acknowledgements

We are grateful to David Ornelles and Michael Imperiale for the 52K mutant viruses, Patrick Hearing for antibodies, Lucy Fisher for help with electron microscopy and the UZH Center for Microscopy and Image Analysis for access to microscopes and virtual machines. We thank Ivo Sbalzarini and all members of the Greber lab for discussions and Maarit Soumalainen for comments on the manuscript. The work was supported by grants from the Swiss National Science Foundation (Grant No. 310030_212802 to UFG), the Natural Sciences and Engineering Research Council of Canada (Grant No. 05048 to NWL), and the Kanton Zürich.

## Authors contributions

Conceptualization: UFG, MB, AGG; methodology: AGG, MB, NWL; investigation: AGG, MB, PB; formal analysis: AGG, PB, MB; data curation: AGG, PB, MB; visualization: AGG, PB; software: AGG, PB; validation: AGG; resources: MOL, NWL; funding acquisition: UFG, NWL; project administration: UFG; supervision: UFG; writing: AGG, UFG; review & editing: AGG, UFG, and input from all authors.

