## Supplemental figure legends for "Spatiotemporal visualization of DNA replication by click chemistry reveals bubbling of viral DNA in virion formation"

### Figure S1: Vinyl-modified nucleosides are incorporated in replicating AdV-C5 and HSV-1 vDNA and can be visualized throughout infection (related to Fig. 1)

(A) Quantitative analyses of vinyl-modified nucleoside incorporation in A549 cells infected with AdV-C5 (MOI 1.5) and clicked with AO-6MT. Nuclei were segmented based on DAPI nuclear signal and the Acridine Orange signal was quantitated in the nuclear area. Data represent means ± SD.

(B) VdU incorporation into HSV-1 replication centers. A549 cells where inoculated with HSV-1 at MOI > 1 at 37°C for 60 min, washed and incubated with VdU until fixation at 8 hpi. Samples were immuno-stained for ICP8 and clicked accordingly. Scale bar, 10 µm.

(C) Quantitative analyses of vinyl-modified nucleoside incorporation in A549 cells infected with HSV-1 (MOI >1) and clicked with AO-6MT. Nuclei were segmented based on DAPI nuclear signal and the Acridine Orange signal was quantitated in the nuclear area. Data represent means ± SD.

(D) Comparative analysis of tetrazine coupling to VdU-labeled DNA. A549 cells where labeled with VdU for 24 h, fixed and clicked with either XFD488-6-MT or 5-TAMRA-6-MT. Dashed lines indicate nuclear outlines. Scale bar, 10 µm.

(E, F) Quantitative analyses of EdC incorporation into HEK-293 and HER-911 cells. Samples were prepared as described in Fig. 1D) and E). Data represent means ± SD of the $N_3$-AlexaFluor488 signal over the DAPI-stained nuclei. Statistical significance was determined by non-parametric ANOVA with Holm-Sidak for multiple comparisons. **, $p<0.0021$; ***, $p<0.0002$; ns, non-significant.

(G) Electron micrograph of negatively stained VdU-labeled AdV-C5_ΔE1 particles. Scale bar, 100 nm.

(H) Representative images and quantitative analysis of AdV-C5_ΔE1-VdU particles stained with AO-6MT. Virions were bound to poly-lysine-coated coverslips, fixed, stained with the anti-hexon 9C12 antibody and AO-6MT clicked. Data represent means of the AO-6MT signal in virus particles. Scale bar, 10 µm.

(I) AdV-C5 progeny production in presence of vinyl-modified nucleosides. A549 cells were infected with AdV-C5 at 37°C for 2 h, removed and labeled with vinyl-modified nucleosides at 8 hpi. Progeny virus was collected at 72 hpi and titrated by $TCID_{50}$. Data represent Spearman-Kärber $TCID_{50}$ values ± SD.

(J) HSV-1 progeny production in presence of vinyl-modified nucleosides. A549 cells were infected with HSV-1 at 37°C for 2 h, removed and labeled with vinyl-modified nucleosides at 2 hpi. Progeny virus was collected at 48 hpi and titrated by $TCID_{50}$. Data represent Spearman-Kärber $TCID_{50}$ values ± SD.

(K) HSV-1-VdU particles stained with AO-6MT. Virions were bound to poly-lysine-coated coverslips, fixed, stained with the anti-HC antibody and AO-6MT clicked. Arrowheads indicate VdU-positive HSV-1 particles. Scale bar, 10 µm.

(L) Tracking of incoming VdU-labeled HSV-1 in cells. A549 cells were inoculated with VdU-labeled HSV-1 for 30 min, washed, incubated 150 min, fixed, stained with anti-HC antibodies and clicked with AO-6MT. Arrowheads indicate VdU-positive HSV-1 particles. Data represent normalized intensities of capsid and vDNA across a dotted line. Scale bar, 10 µm.

### Figure S2: VdU-AO-6MT labels viral and host DNA, stalls viral replication and resolves DNA rearrangement in infection (related to Fig. 2)

(A, B) Comparative analysis of EdC and VdU incorporation in AdV-C5 infection. A549 cells were infected with AdV-C5 (MOI 3) for 60 min, washed, pulsed with 2.5 µM EdC at 16-20 hpi, pulsed with 50 µM VdU at 20-24 hpi (B) or vice versa (C) and fixed at 24 hpi. Samples were stained with anti-DBP antibodies (cyan), clicked with AO-6MT (magenta) and clicked with $N_3$-AlexaFluor647 (green). Scale bar, 10 µM.

(C) Quantification of EdC-labeled vDNA overlap with DBP. Samples were prepared as described in Fig. 2B. EdC-labeled vDNA structures were 3D-segmented based on the EdC-Alexa647 signal. Overlap was computed against the 3D-segmented DBP structures. Each data point represents an EdC-labeled vDNA structure. Median is indicated in red.

(D) Quantitative analysis of DBP overlap with VdU-labeled vDNA. Samples were prepared as described in Fig. 2B. DBP structures were 3D-segmented based on the DBP signal. Overlap was computed against the 3D-segmented VdU-labeled vDNA structures. Each data point represents a DBP structure. Median is indicated in red.

(E) Quantitative analysis of VdU-labeled vDNA overlap with DBP. Samples were prepared as described in Fig. 2B. VdU structures were 3D-segmented based on the VdU-AO-6MT signal. Overlap was computed against the 3D-segmented DBP structures. Each data point represents a VdU-labeled vDNA structure. Median is indicated in red.

(F) Dose-dependent incorporation of EdC during AdV-C5 infection in VdU-AO-6MT DNA labeled cells. A549 cells were grown in different concentrations of VdU, followed by infection with AdV-C5 (MOI 3) for 60 min, washed and labeled with AO-6MT (concentrations adjusted to the VdU concentrations) and 2.5 µM EdC in FluoroBrite at 6-44 hpi. Cells were fixed at 44 hpi, clicked with $N_3$-AlexaFluor647 and stained with DAPI. Data represent the integrated VdU-AO-6MT and EdC-AlexaFluor647 intensities measured over the DAPI-stained nuclei. Each data point represents one nucleus.

(G) Comparative analysis of the cellular and viral DNA distribution in AdV-C5 infection. Samples were prepared as described in the previous panel. Dashed lines indicate nuclear outlines. Zoomed nuclei show EdC signal localizing at the nuclei periphery. Data represent EdC-labeled vDNA, VdU-labeled cell DNA and DAPI normalized intensities measured over the white dotted line. Arrows indicate chromatin condensation events of viral origin. Images are maximum projections. Scale bar, 10 µm.

(H, I) Live cell imaging of VdU-AO-6MT labeled host DNA during AdV-C5 infection, including the 44 hpi endpoint. Samples were prepared as described in panel F). Live imaging was from 8-44 hpi at a frequency of four frames per hour. Arrowheads indicate cellular chromatin condensation events. The prominent extranuclear signals represent acidic compartments, where AO-6MT tends to accumulate over time. Images shown are maximum projections. Scale bar, 10 µm.

(J) Changes in nuclear morphology during AdV-C5 infection. A549 cells were infected with AdV-C5 (MOI 3) for 60 min, washed, fixed at the indicated times pi and stained with DAPI.

Data represent the nuclear area and the DAPI mean intensity within each nuclei. Scale bar, 10 µm.

(K) Live cell imaging of AdV-C5 VRCs. A549 cells were inoculated with AdV-C5 (MOI 3) for 60 min, washed and incubated with 25 µM VdU from 18-22 hpi, followed by AO-6MT (10 µM) addition in FluoroBrite containing Hoechst DNA dye. Live imaging was from 23-65 hpi at a frequency of two frames per hour. Images shown are maximum projections. Scale bar, 50 µm.

(L) Quantitative analysis of EdC incorporation in VdU-AO-6MT labeled AdV-C5 infection. A549 cells were infected with AdV-C5 (MOI 3) for 120min, washed, pulse-labeled with VdU at 16-20 hpi, clicked with AO-6MT at 20-24 hpi and labeled with 2.5 µM EdC at 24-28 hpi. Cells were fixed at 28 hpi, clicked with $N_3$-AlexaFluor647 and stained with DAPI. Data represents percentage of double nucleoside-positive DAPI-stained nuclei across various VdU and AO-6MT concentrations.

(M) Live imaging of HSV-1 VRCs. A549 cells were inoculated with HSV-1 (MOI > 1) for 60 min, washed and incubated with 25 µM VdU from 1-6 hpi, followed by AO-6MT (10 µM) in FluoroBrite containing Hoechst at 7 hpi. Live imaging was from 8-40 hpi as described in panel K). Images are maximum projections. Scale bar, 50 µm.

(N) Quantitative analysis of EdC incorporation in VdU-AO-6MT labeled HSV-1 infection. A549 cells were infected with HSV-1 (MOI > 1) for 60 min, washed, pulse-labeled with VdU at 1-8 hpi, clicked with AO-6MT at 8-12 hpi and labeled with 2.5 µM EdC at 12-16 hpi. Cells were fixed at 16 hpi, clicked with $N_3$-AlexaFluor647 and stained with DAPI. Data represents percentage of double nucleoside-positive DAPI-stained nuclei across various VdU and AO-6MT concentrations.

### Figure S3: Late in infection, early-replicated vDNA dislocates from the VRC and does not contain the packaging factor IVa2 unlike late-replicated vDNA in the VRC (related to Fig. 3)

Representative images and quantitative analysis of differentially-labeled vDNA co-localization with AdV-C5 proteins IIIa (A), 52K (B), IVa2 (C), 33K (D), V(E) and VII (F) at late stages of infection (34 hpi). Specimens were labeled with 2.5 µM EdC at 12-14 hpi and with 50 µM VdU at 32-34 hpi, stained with diverse antibodies (cyan), and clicked with AO-6MT (magenta) and $N_3$-AlexaFluor594 (green). Each data point represents a vDNA structure. Median is indicated in red. Images are maximum projections. White squares show regions at higher contrast to highlight vDNA puncta. Arrowheads indicate single VdU-labeled puncta (labeled at 32 hpi). Scale bar, 10 µm.

### Figure S4: Viral DNA puncta do not co-localize with DBP or the packaging factors during late stages of infection (related to Fig. 4)

(A) Representative image of EdC-labeled vDNA puncta in A549 nuclei upon cell infection with AdV-C5 wt for 60 min, washing, pulse-labeling with 2.5 µM EdC at 22-24 hpi, pulse-labeling with 50 µM VdU at 32-34 hpi and fixation at 34 hpi. Samples were clicked with AO-6MT (magenta) and $N_3$-AlexaFluor594 (green). Images are maximum projections. Scale bar, 10 µm.

(B) Quantitative analysis of co-localization of the different vDNA species co-localization with DBP at late stages of infection. Samples were prepared as described in Fig. 4B. Data distributions are shown as violin plots. Median is indicated in red. Statistical significance was determined by non-parametric ANOVA with Holm-Sidak for multiple comparisons. ***, $p<0.0002$.

(C-F) Representative images and quantitative analysis of differentially-labeled vDNA puncta co-localization with AdV packaging proteins in late stages of infection. Samples were prepared as described in Fig. 4B. Data distributions are shown as violin plots. Median is indicated in red. Statistical significance was determined as indicated in B). *, $p<0.03$; **, $p<0.0021$; ***, $p<0.0002$; ****, $p<0.0001$; ns, non-significant. Arrows indicate single EdC-puncta (labeled at 22 hpi); arrowheads indicate single VdU-puncta (labeled at 32 hpi). Images are maximum projections. Scale bar, 10 µm.

**Figure S5: The formation of vDNA structures containing GFV-V is replication dependent (related to Fig. 5)**

(A) Comparative analysis of GFP-V, but not anti-V, co-localization with EdC-labeled vDNA puncta. Samples were prepared as described in Fig. 5A. Arrows indicate GFP-V-positive, but not anti-V-positive EdC-labeled vDNA puncta. Data represent normalized intensities of vDNA (magenta), GFP-V (green) and anti-V (cyan) across a dotted line. Scale bar, 5 µm.

(B) Representative images of background signals for AdV-C5 core proteins V and VII. Fixed, non-infected A549 cells were stained with anti-V or anti-VII and DAPI. Dashed lines indicate nuclear outlines. Images are maximum projections. Scale bar, 10 µm.

(C) Live imaging of replication-dependent AdV-C5_GFP-V viral core morphogenesis. A549 cells were infected with AdV-C2-GFP-V as described in Fig. 5A. DFT was added at 24 hpi and kept for the rest of the infection. GFP-V signal was recorded on full cell Z-stacks from 26-27 hpi at a frequency of four stacks per minute. Arrowheads indicate viral core nucleation events. Images are single Z slices. Scale bar, 1 µm.

**Movie S1: Cellular DNA clustering during Adenovirus infection**

A549 cells were grown in 25 µM VdU. Cells were infected with AdV-C5 (MOI 3) for 60 min, washed and labelled with 10 µM AO-6MT in FluoroBrite at 6-44 hpi. AO-6MT was recorded from 8-44 hpi at a frequency of four frames per hour in an IXMc confocal spinning-disk microscope. Timestamps are in h:min. The prominent extranuclear signals represent acidic compartments, where AO-6MT tends to accumulate over time. Frames are maximum projections.

**Movie S2: AdV-C5 VRCs resolved by AO-6MT**

A549 cells were inoculated with AdV-C5 (MOI 3) for 60 min, washed and incubated with 25 µM VdU from 18-22 hpi, followed by 10 µM AO-6MT in FluoroBrite containing Hoechst DNA dye. AO-6MT was recorded from 23-65 hpi at a frequency of two frames per hour in an IXMc confocal spinning-disk microscope. Timestamps are in h:min. Frames are maximum projections.

#### Movie S3: HSV-1 VRCs resolved by AO-6MT

A549 cells were inoculated with HSV-1 (MOI > 1) for 60 min, washed and incubated with 25 µM VdU from 1-6 hpi, followed by 10 µM AO-6MT in FluoroBrite containing Hoechst at 7 hpi. AO-6MT was recorded from 8-40 hpi as described in movie S2. Timestamps are in h:min. Frames are maximum projections.

#### Movie S4: GFP-V bubbling from the VRC during late stages of AdV infection

A549 cells were infected with AdV-C2-GFP-V for 60min, washed, pulse-labeled with 2.5 µM EdC at 22-24 hpi. GFP-V was recorded on full cell Z-stacks from 27:30-28:00 hpi at a frequency of four stacks per minute in an Olympus IXplore SpinSR10 spinning-disk microscope. Timestamps are in h:min:sec. Frames are single Z slices.

#### Movie S5: Zoomed section of a GFP-V-bubbling active zone

A549 cells were infected and imaged as described in movie S4. Timestamps are in h:min:sec. Frames are single Z slices.

#### Movie S6: Frequency and motility reduction of GFP-V bubbling events in replication-inhibited AdV infection

A549 cells were infected with AdV-C2-GFP-V as described movie S4. DFT (2.5 µM) was added at 24 hpi and kept for the rest of the infection. GFP-V signal was recorded on full cell Z-stacks from 26-27 hpi at a frequency of four stacks per minute in an Olympus IXplore SpinSR10 spinning-disk microscope. Arrowheads indicate viral core nucleation events. Timestamps are in h:min:sec. Frames are single Z slices.

**Movie S7: Zoomed section of a GFP-V-bubbling event in replication-inhibited AdV infection**

A549 cells were infected and imaged as described in movie S6. Timestamps are in h:min:sec. Frames are single Z slices.
