## Supplementary figures and images for "Spatiotemporal visualization of DNA replication by click chemistry reveals bubbling of viral DNA in virion formation"

### Supplemental figure 1

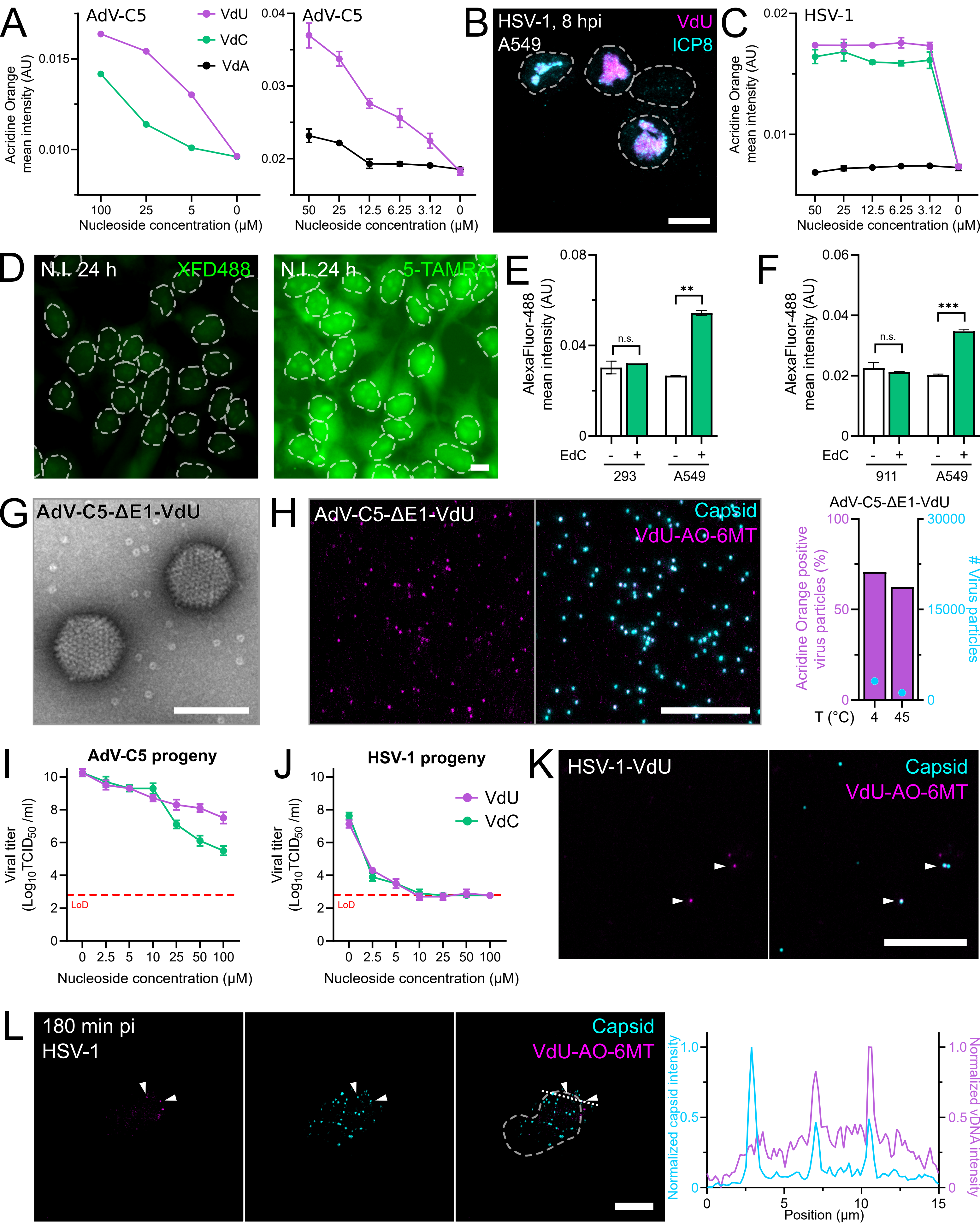

### Supplemental figure 2

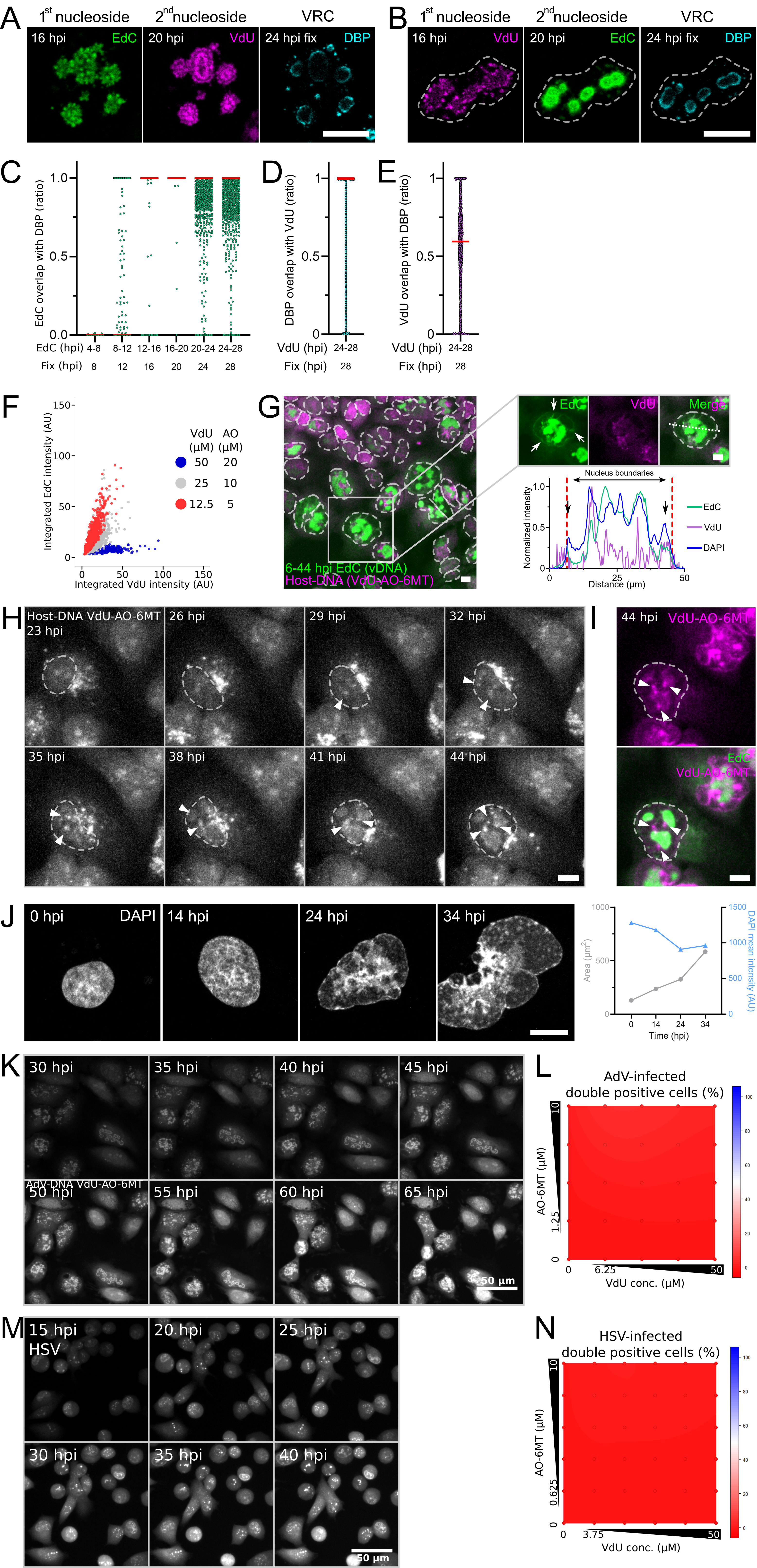

### Supplemental figure 3

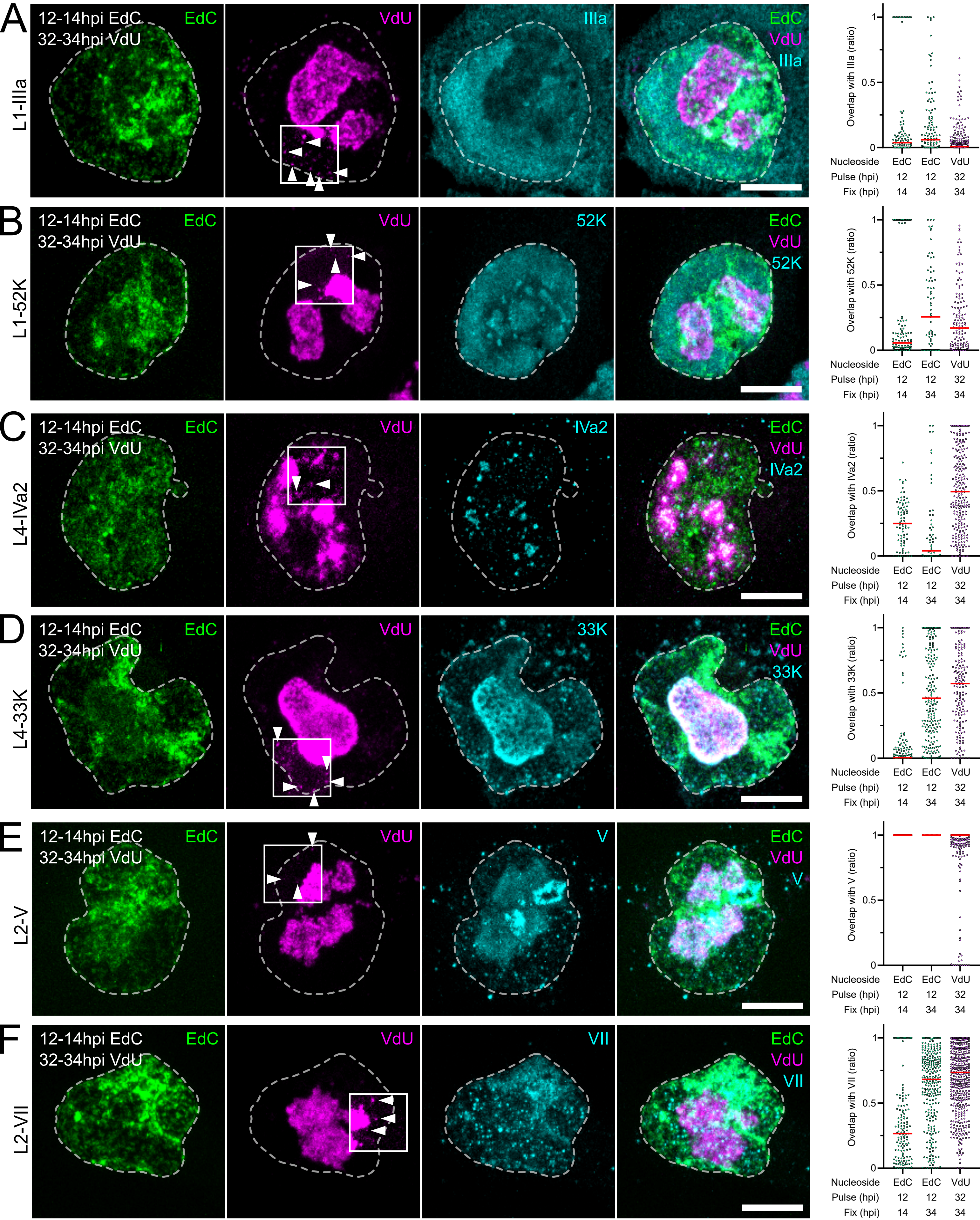

### Supplemental figure 4

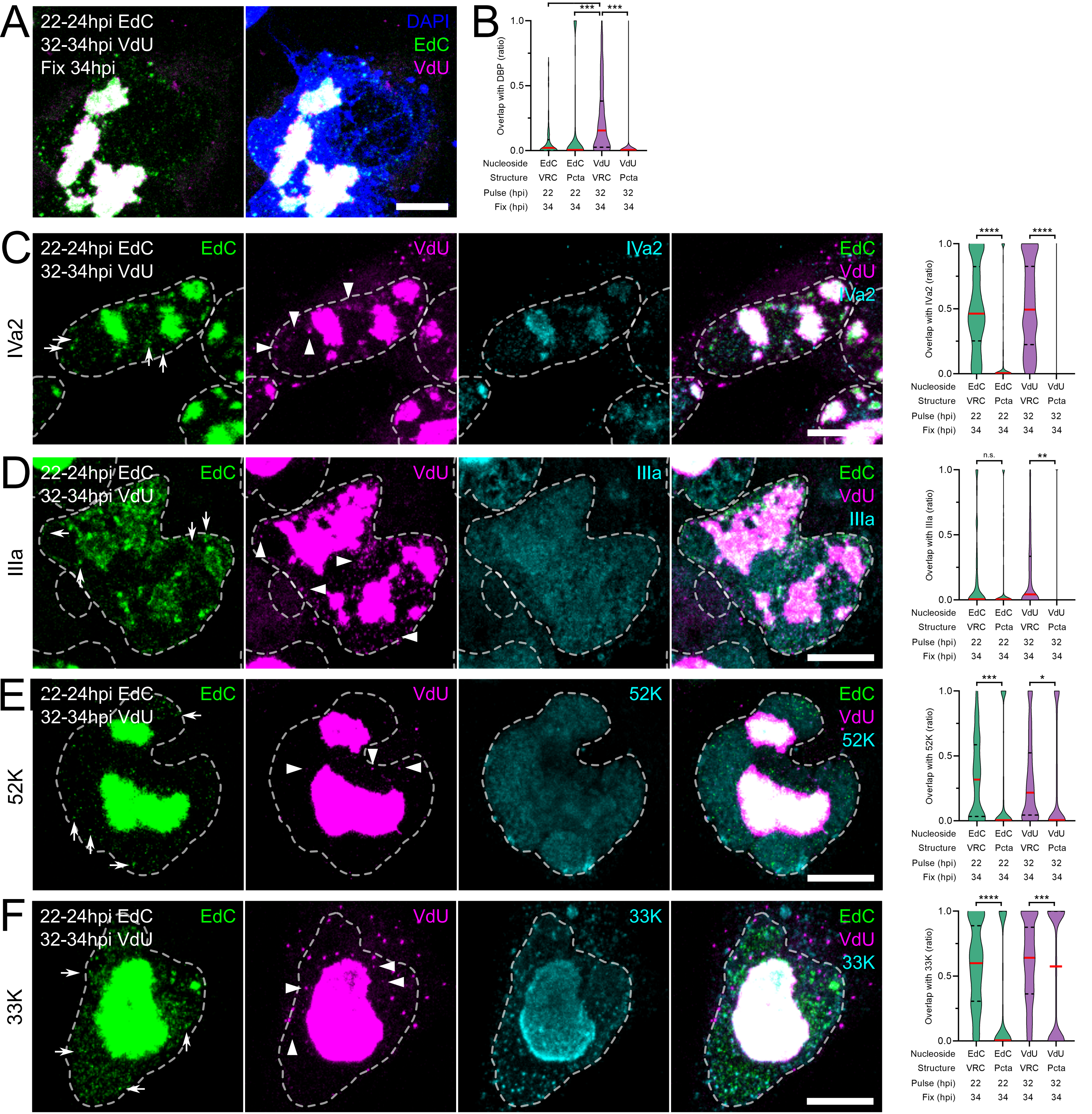

### Supplemental figure 5

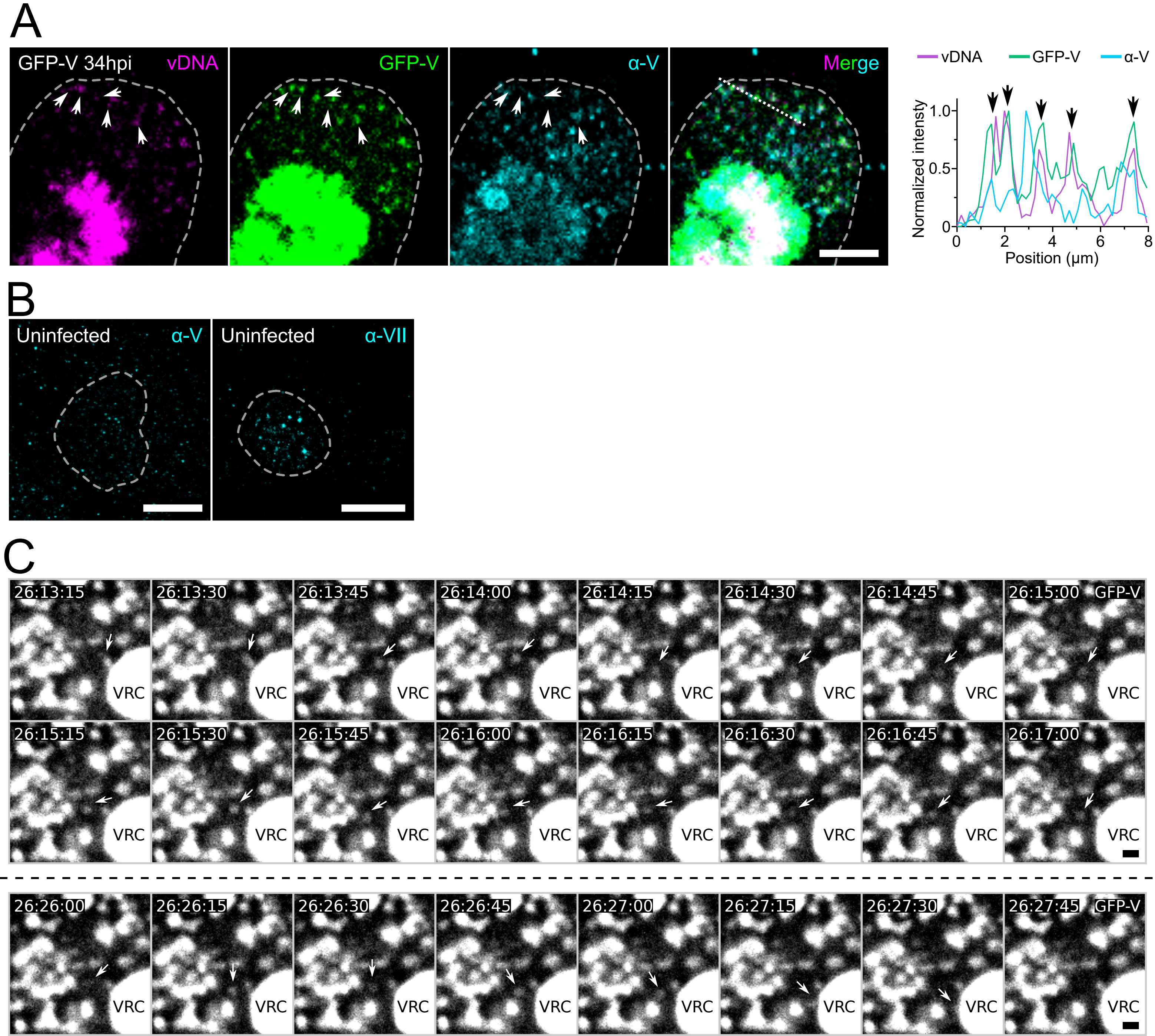
